# Systematic and dynamic task coding in the frontal cortex

**DOI:** 10.64898/2026.11.10.765810

**Authors:** JeongJun Park, Hyung-Bum Park, Charles D. Holmes, Lawrence H. Snyder

## Abstract

Behavioral flexibility depends on the capacity to form and utilize task representations. Prior work has shown that baseline activity in the prefrontal cortex (PFC) can differ between two tasks. Comparing more than two tasks makes it possible to ask whether different neuronal populations share a common form of task encoding, that is, a conserved representational geometry across tasks. We trained two rhesus monkeys to perform 6 cognitive tasks and recorded spiking activity simultaneously from hundreds of neurons in the frontal eye field (FEF) or the dorsolateral PFC (dlPFC), one area per session. During the fixation period, before any task-specific stimulus appeared, the population activity state already reflected the upcoming task, and task identity could be decoded on single trials. Strikingly, the geometry of the 6 task states was conserved across populations recorded in different sessions, across the two areas, and across the two monkeys. Global signals that differ between tasks, such as expected reward, task difficulty, and attention, measured by reward amount, hit rate, and reaction time, did not fully account for this shared geometry. These findings suggest that the frontal cortex systematically adjusts its population state to encode and prepare for the upcoming task.

## Introduction

Primates, including humans, can rapidly switch between previously learned tasks. How the brain supports such task switching remains an open question. One influential framework for understanding this function is the concept of a “task set”: an abstract configuration of perceptual, attentional, mnemonic, and motor processes that is maintained to guide the current task^1^. A task set is distinct from long-term task knowledge or schema, which stores the structure of a task but does not by itself guide action^2^. Lateral prefrontal cortex (LPFC) has been proposed to construct task sets by reformatting this stored knowledge into active control representations that directly support action selection, transforming schematic knowledge into executable policies^3^. Because task sets can be established prior to the start of a task, they could be instantiated as preparatory activity states that allow for faster and more efficient task-related information processing^4,5^.

Consistent with this view, previous studies reported that failures in the preparatory process can lead to deficits in task performance, particularly following PFC damage^6^. Although baseline or preparatory activity in the PFC can differ across tasks or rules^7–9^, the neural representations and mechanisms underlying task preparatory states remain poorly understood. Most neuroimaging and neurophysiological studies of task coding in the PFC have focused on within-trial dynamics, examining how neurons represent stimuli, decisions, or memory contents during task execution^10–13^. Those studies that considered the preparatory period used only a limited number of tasks (often just two) and lacked population-level recordings.

To address how the PFC represents task context in its preparatory population activity, we trained two rhesus monkeys to perform 6 tasks that shared some features but differed in others. Comparing 6 tasks, rather than two, allowed us to ask whether different neuronal populations share a common geometry of task encoding. In each session, we simultaneously recorded the spiking activity of hundreds of neurons in either the frontal eye field (FEF) or the dorsolateral PFC (dlPFC). Tasks were presented in blocks, and each trial included a short fixation interval, before any task-specific stimulus appeared, during which the animals could prepare for the upcoming trial. Using a neural geometry approach, we found that population activity during this interval already reflected the upcoming task, and task identity could be decoded on single trials. These task-specific preparatory states could not be explained by the previous task or the task cue alone, and they weakened on error trials. Strikingly, their geometry was conserved across sessions, areas, and animals, and was not fully explained by global signals such as reward. These findings suggest that the frontal cortex systematically adjusts its population state to encode and prepare for the upcoming task.

## Results

### Six cognitive tasks with shared and differing features

To examine whether PFC population activity distinguishes multiple task conditions, we trained two monkeys to perform six different visual tasks that share some features but differ in others (Figure 1A–F, see legend for details). Each trial consisted of a 1 s fixation period followed by several task-relevant stimuli, and a final saccade decision based on the rules of the current task. To allow animals to anticipate the upcoming task before task-specific targets appeared, we used a block design in which each block contained 16 successful trials of the same task, with each task associated with a different fixation cross color (except that the two non-match-to-sample tasks shared the same color). To control for the color effect and isolate task-related signals in the preparatory neural activity, the fixation cross and spatial targets were presented in white during the last 8 trials of each 16-trial block. We collected about 24 blocks per task in each of 32 sessions (23.6 ± 2.4 blocks per task and 2265 ± 229 correct trials per session, mean ± SD).

**Figure 1.**
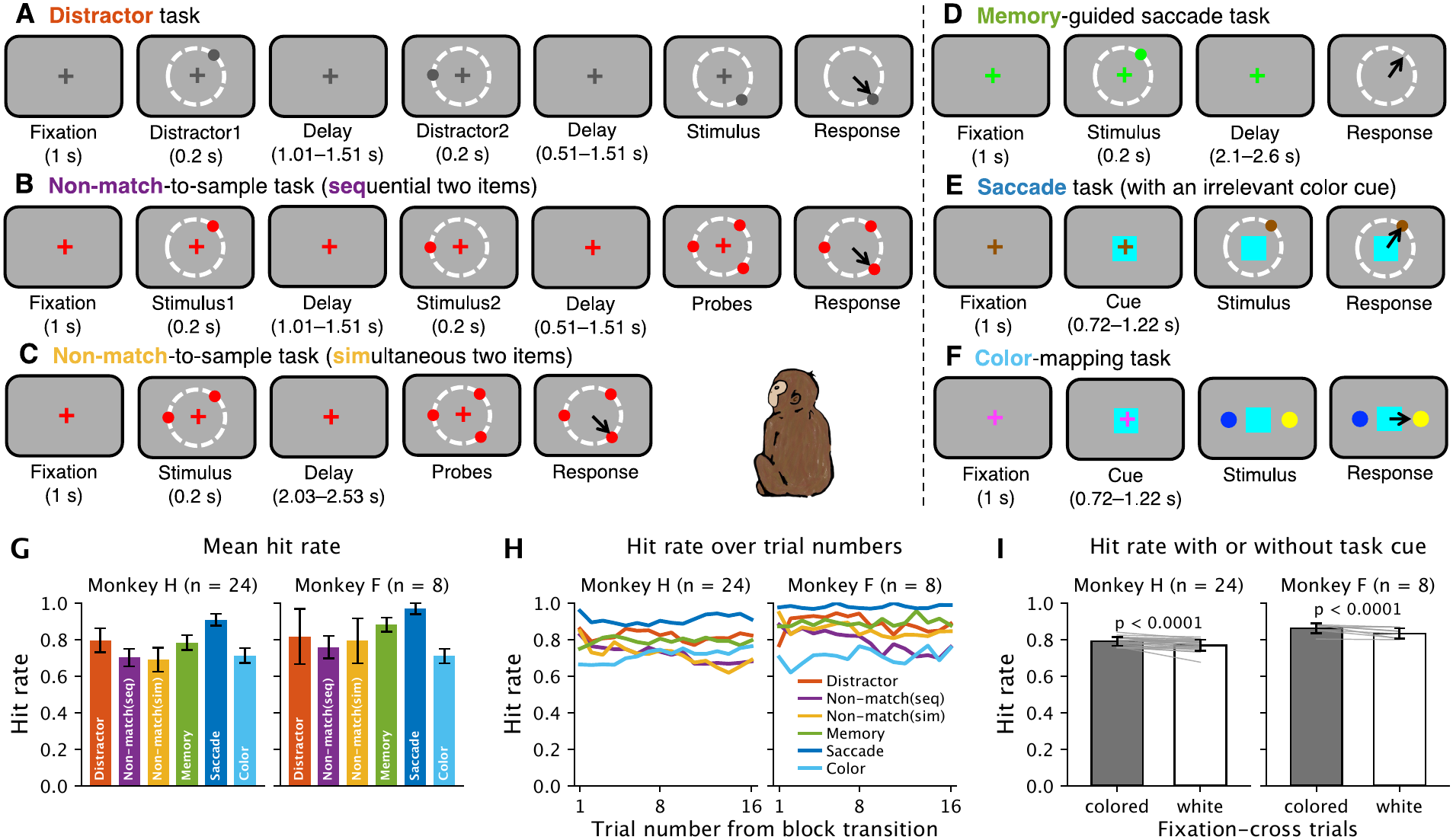
Task design and behavioral performance. Each trial begins with a 1-second fixation period at a central cross whose color indicates task identity (A: grey, B–C: red, D: green, E: brown, F: magenta). In tasks with peripheral spatial targets (A–E), target colors match the fixation cross color. Tasks are interleaved in 16-trial blocks, with the fixation cross and peripheral spatial targets turning white during the final 8 trials of each block across all tasks. In each task, monkeys are rewarded for making the appropriate saccade when cued to move by the extinction of the fixation cross. In five tasks (Figure 1A–E), the spatial location(s) of the target(s) are task-relevant, whereas in the color-mapping task (Figure 1F) the color is relevant. Three of the spatial tasks (Figure 1B–D) require the target location(s) to be held in working memory, whereas the other two (Figure 1A, E) do not. The distractor task (Figure 1A) is identical to the sequential non-match-to-sample task (Figure 1B) except that only a single (novel) target appears after the delay. The memory-guided saccade task (Figure 1D) and the saccade task (Figure 1E) share a single spatial target that disappears prior to the go cue in the memory task. A colored square appears behind the fixation cross in both the saccade task (Figure 1E) and the color-mapping task (Figure 1F), but the color is task-relevant only in the color-mapping task. **(A)** Distractor task. Two distractor targets are presented sequentially before the saccade target. The animal must ignore the first two targets and make a saccade to the third target upon disappearance of the fixation cross. **(B–C)** Non-match-to-sample task with two memory targets presented sequentially (B) or simultaneously (C). After the final delay, the memory targets reappear along with a novel target. The animal must saccade to the novel target. **(D)** Memory-guided saccade task. After a delay period, the animal must make a saccade to the target location. **(E)** Saccade task with an irrelevant color cue. The animal must make a saccade to a peripheral target in the presence of a task-irrelevant colored square (cyan, blue, or yellow). **(F)** Color-mapping task. The animal was trained to associate arbitrary cue–target color pairs (Monkey F: blue→cyan, cyan→yellow, yellow→blue, Monkey H: blue→blue, cyan→cyan, yellow→yellow). Based on the color of the central cue, the animal makes a saccade to the corresponding target color among two peripheral stimuli, irrespective of spatial location. **(G)** Hit rate for each task (mean ± SD across 24 sessions in Monkey H and 8 sessions in Monkey F). Chance is 33% in B and C and 50% in F. In A and E, single target is on the screen at the go cue. In D, a saccade of the appropriate amplitude in a random direction would land within the response window (radius 8.5 dva) on 21% of trials. **(H)** Hit rate for each task as a function of trial number from the block transition. Trial number counts all trials. The fixation cross turns white after the 8th successful trial. **(I)** Hit rate (as in H) with or without task cue: trials 1 through 8 (with a task-colored fixation cross) versus 9 through 16 (white fixation cross). Bars: mean ± SD across sessions. Lines: individual sessions.

Monkeys performed well in all tasks, with hit rates ranging from 69–91% across tasks in Monkey H and 71–97% in Monkey F (Figure 1G). We asked whether animals recognized task transitions immediately or required several trials to adjust their behavior. Performance on the first trial after a block transition was not lower than on the rest of the block (Figure 1H). The first trial was even slightly more accurate than trials 2 to 16 in Monkey H (82% versus 78%, trial-label permutation test of the trial position within each block, p < 0.0001), but not in Monkey F (86% versus 85%, p = 0.53). This indicates immediate behavioral adjustment. To test whether animals relied on the task cue or also used the block structure and other cues to track the current task, we examined performance after the fixation cross and spatial target colors changed to white following the 8th successful trial. Hit rate dropped but only slightly (Figure 1I, from 79% to 77% in Monkey H and from 86% to 83% in Monkey F, trial-label permutation test of the cue stage within each block, p < 0.0001 in both monkeys).

### Representational geometry of task contexts before task onset

We hypothesized that information about the upcoming task might systematically modulate PFC baseline activity^7–9^. Using a Neuropixels 1.0 NHP probe, we recorded spiking activity from hundreds of neurons simultaneously in the frontal eye field (FEF) or the dorsolateral prefrontal cortex (dlPFC) while monkeys performed six tasks (Supplementary Figure 1). In total, we recorded 8,934 neurons in Monkey H (6,419 from 13 FEF sessions and 2,515 from 11 dlPFC sessions) and 2,559 neurons in Monkey F (1,667 from 5 FEF sessions and 892 from 3 dlPFC sessions). We analyzed the fixation period, before any task-specific stimulus appeared.

We first tested whether task identity was encoded in the mean firing rate across the entire population. One can imagine that differential engagement across tasks might lead to higher baseline activity in some tasks compared to others. This was not the case: during the initial fixation period, prior to receipt of any task-relevant stimuli, the mean firing rate largely did not differ among tasks in both areas of both monkeys (Figure 2A). We next tested whether the activity pattern might be differentially modulated across tasks. At the individual cell level, 15% of FEF cells and 17% of dlPFC cells in Monkey H and 10% of FEF cells and 6% of dlPFC cells in Monkey F differed in fixation-period firing rate among tasks (one-way ANOVA of block-mean rates, p < 0.01). To look for patterns across the entire population, we used linear discriminant analysis (LDA) to define a low-dimensional neural subspace spanned by the components that best separated the six tasks (which we refer to as “task components”) and projected population activity onto this subspace (which we termed “task subspace”). To ensure that we were identifying a reproducible signal and not just fitting noise, the components were estimated from training blocks and then held-out blocks or trials were projected onto those components (see Materials and Methods). In this task subspace, the six tasks occupied distinct positions, except for the two non-match tasks, which largely overlapped (Figure 2B). We held out the blocks rather than trials because population activity drifted slowly over time, so that trials within a block resembled each other more than trials of other blocks of the same task (Supplementary Figure 2A–B). Holding out single trials instead would have let this block-specific signal leak into the training set (Supplementary Figure 2C).

**Figure 2.**
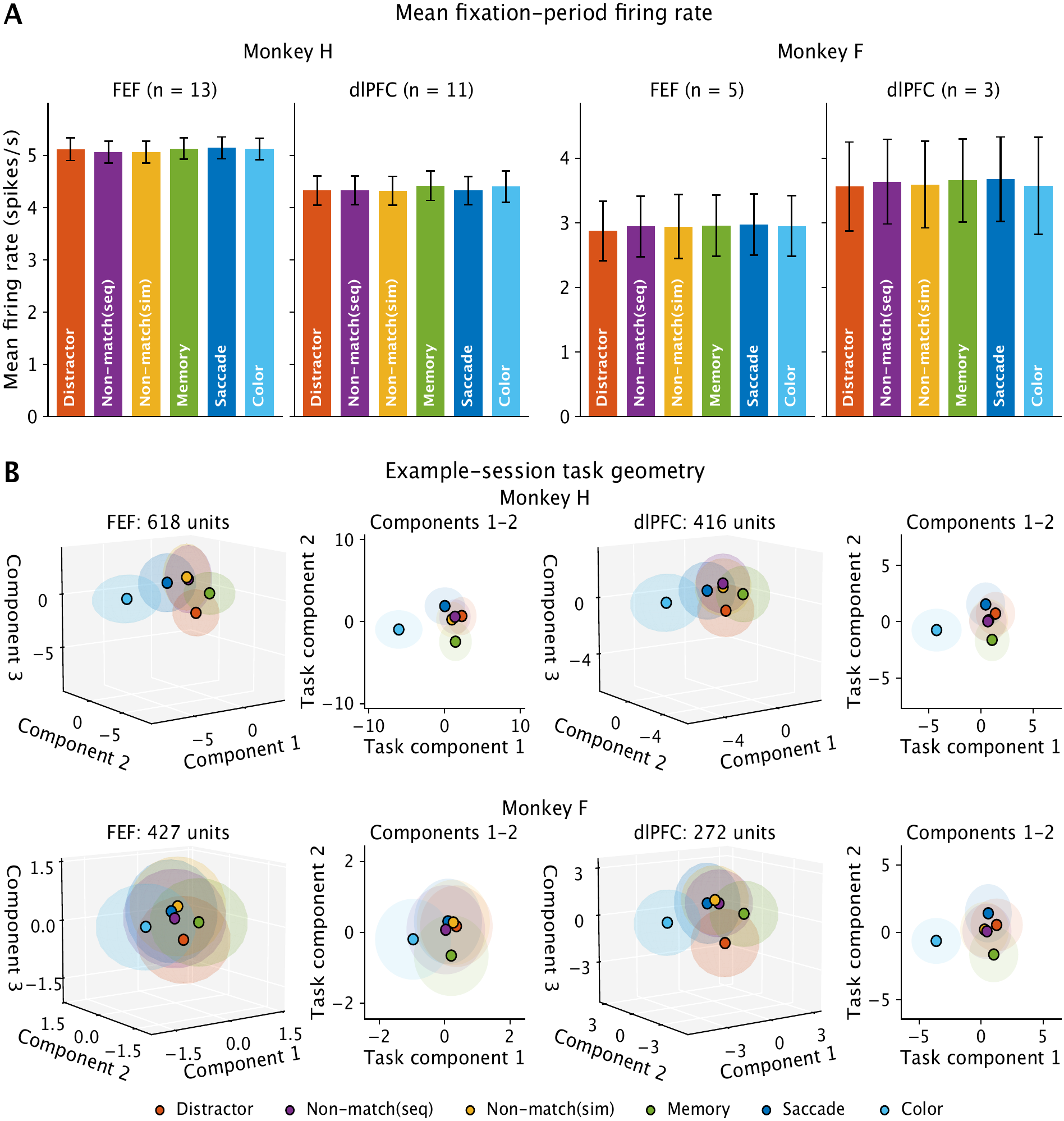
Mean firing rate is task-independent, but the population state encodes the task. **(A)** Mean fixation-period firing rate for each task in the FEF and dlPFC sessions of each monkey (mean ± SEM across sessions, 13 FEF and 11 dlPFC sessions in Monkey H and 5 FEF and 3 dlPFC sessions in Monkey F). The only statistically detectable difference among tasks, in the FEF of Monkey H, was at most 0.09 spikes/s (Friedman test across sessions, p = 0.003, versus p > 0.15 elsewhere). **(B)** Task geometry in one example FEF session and one example dlPFC session per monkey. In each session, the task components were estimated from training blocks, and the held-out blocks and successful trials were projected onto those estimated components. The configuration of the six task centroids in the first four components was rotated by generalized Procrustes alignment method such that all panels share one orientation (see Materials and Methods). The resulting task configurations are shown on the first three aligned axes (left) and the first two (right). Dots: mean of each task across blocks. Shading: ±1 SD across single trials of that task. Axes are in units of z-scored firing rate. The number of units is given above each panel.

### Task identity can be decoded from fixation-period activity only in successful trials

To quantify this representation, we decoded task identity from held-out neural data by projecting population activity patterns into the task subspace and assigning each pattern to the task whose centroid in the training blocks was nearest. Decoding the block-averaged activity of held-out blocks was far above chance (16.7%) in every session (Figure 3A–B). With the first four task components, accuracy was 78 ± 6% correct in FEF and 65 ± 12% in dlPFC of Monkey H and 53 ± 17% and 67 ± 14% in Monkey F (mean ± SD across sessions, block-label permutation test of the task within each session, p < 0.0001 for every number of components). Task identity was also decodable from the fixation-period activity of single trials (Figure 3C–D, first four components, 47 ± 3% in FEF and 37 ± 7% in dlPFC of Monkey H and 31 ± 9% and 38 ± 11% in Monkey F, mean ± SD, p < 0.0001 for every number of components). With the first four components, both block and single-trial decoding were above chance in every individual session (block-label permutation test of the task within each session, p < 0.05). Accuracy increased as components were added up to the fourth (Figure 3A and C, brackets, one-sided test of the gain from adding a component). In Monkey H, every step from one to four components increased accuracy (p < 0.0001). In Monkey F, block and single-trial decoding accuracy increased significantly at every step from 1 to 4 components (p < 0.005), except the 3→4 step in FEF, which increased significantly for single-trial decoding (p = 0.04) but not for block decoding (p = 0.11). The fifth component did not increase accuracy in either monkey (p > 0.8). We retained a component whenever adding it increased accuracy, even if not significantly (as for the 3→4 step of block decoding in FEF of Monkey F), and used the first four components in all subsequent analyses.

**Figure 3.**
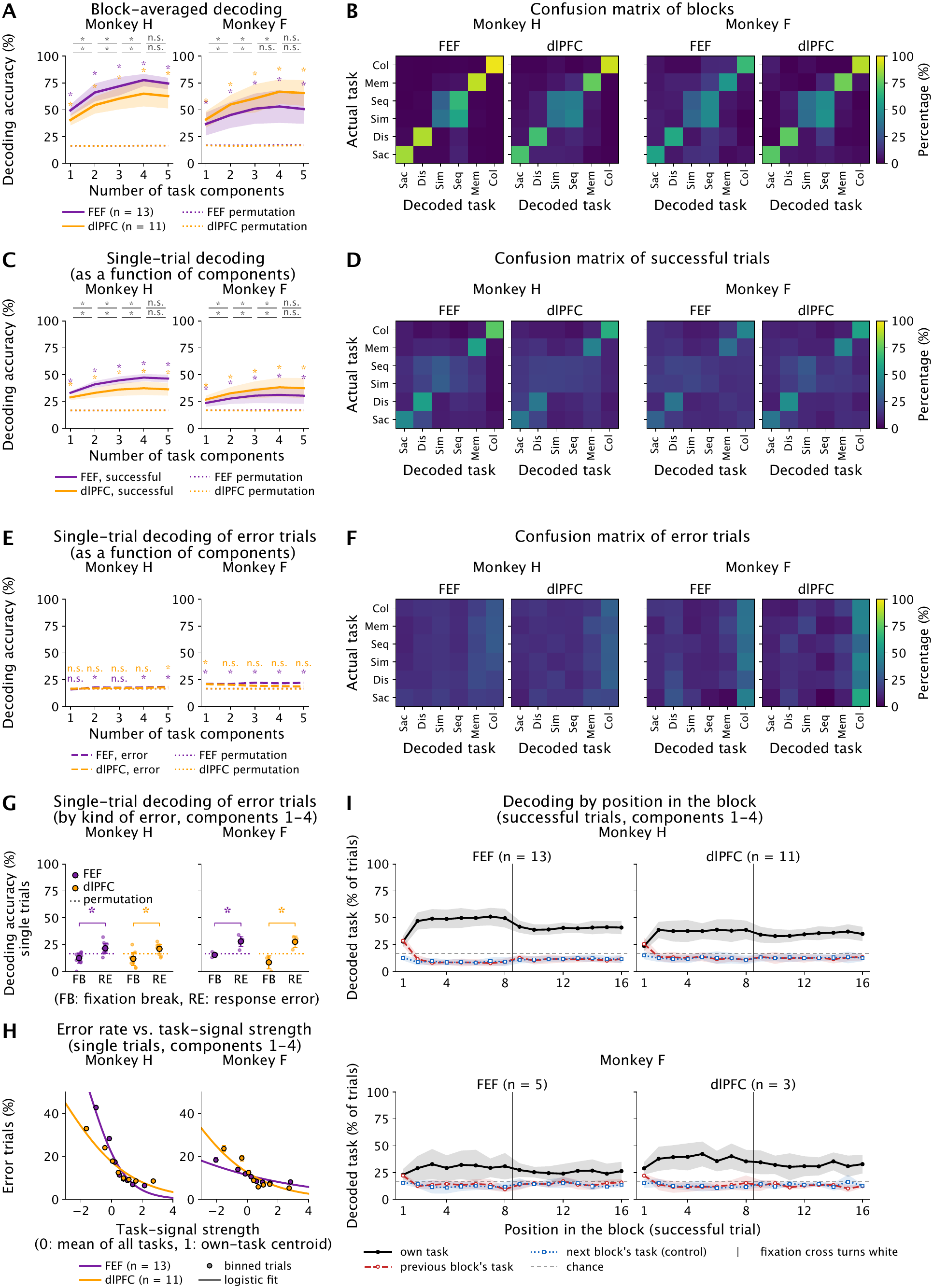
Within-session task decoding from fixation-period activity. Decoding of held-out data by a nearest-centroid classifier in the subspace of the first k task components (13 FEF and 11 dlPFC sessions in Monkey H and 5 FEF and 3 dlPFC sessions in Monkey F). **(A)** Decoding accuracy for the block-averaged activity of held-out blocks as a function of the number of task components (mean ± SD across sessions). Dotted lines, mean accuracy after permuting the task labels across blocks (100 permutations per session). Asterisks above each point, accuracy significantly above this permutation null (block-label permutation test of the task within each session, p < 0.05). Brackets above the panels compare accuracy with k components to accuracy with k − 1 components (upper row, dlPFC, and lower row, FEF). The label marks whether adding component k increased accuracy more than under permutation (one-sided test, asterisk for p < 0.05 and n.s. for not significant). **(B)** Confusion matrices for held-out blocks using first four components: session-averaged percentage of items of each actual task (rows) decoded as each task (columns). Sac (saccade), Dis (distractor), Sim and Seq (non-match-to-sample with simultaneous and sequential items), Mem (memory-guided saccade) and Col (color-mapping). **(C)** Decoding accuracy for single held-out successful trials. Lines and shading: mean ± SD across sessions. Dotted lines, asterisks and brackets are as in A. **(D)** Confusion matrices for held-out successful trials using first four components. **(E)** Decoding accuracy for single held-out error trials, decoded with the decoder trained on successful trials. Dashed lines: mean across sessions. Dotted lines as in A. Asterisks and n.s. above the curves mark whether error-trial accuracy at each k is significantly above the permutation null (lower row, FEF, and upper row, dlPFC, p < 0.05). **(F)** Confusion matrices for held-out error trials using first four components. **(G)** Single-trial decoding accuracy of held-out error trials by error type. FB: fixation break after the fixation period and before the go cue, RE: response error after the go cue. Small dots: sessions. Large dots: mean ± SD across sessions (purple for FEF and orange for dlPFC). Dotted lines: mean accuracy of the error trials after permuting the task labels across blocks. Brackets: fixation break versus response error in each area, trial-label permutation test of the kind of error within each session, sessions combined (asterisk, p < 0.05). **(H)** Probability of an error as a function of the task-signal strength of the trial, that is, its position on the axis from the mean of all six tasks (0) to the centroid of its own task (1) in the subspace of the first four components, for all held-out trials of the sessions of each area (purple for FEF and orange for dlPFC). Dots: error rate of the trials in eight bins of task-signal strength with equal numbers of trials (mean ± SE of a proportion). Lines: logistic regression pooled over the sessions of each area with one intercept per session. The slope differs from the null obtained by shuffling the success and error labels of trials in each session and task in every area (1,000 permutations, p < 0.001). **(I)** Decoding of held-out successful trials as a function of their position in the block (successful trials 1 to 16), with one decoder per fold trained on trials 3 to 16 of the training blocks (first four components). Black solid line: percentage of trials decoded as their own task. Red dashed line: percentage decoded as the task of the previous block. Blue dotted line: percentage decoded as the task of the next block, a control. The previous-and next-block readouts include only blocks that follow or precede a switch to a different task, excluding switches between the two non-match tasks, which share the red fixation cross. Lines and shading: mean ± SD across sessions. Gray dashed line: chance for the own-task readout (16.7%). A trial decoded as its own task cannot also be decoded as the previous or next task, so chance for red and blue lines is below 16.7% and changes with own-task accuracy. The previous-block readout is compared with the next-block readout, which serves as a null because the next task is not predictable (see Materials and Methods). Vertical line: the fixation cross turns white after the 8th successful trial.

Consistent with the geometry, the color-mapping task was decoded most reliably whereas the two non-match tasks were frequently confused with each other (Figure 3B and D). Both block and single-trial decoding accuracy were higher in FEF than in dlPFC in Monkey H, but FEF sessions also yielded about twice as many neurons. When decoding was based on 100 randomly drawn neurons per session and the first four components, the difference in decoding accuracy in Monkey H shrank from 13 to 6 percentage points for block decoding (FEF versus dlPFC, 64 ± 9% versus 58 ± 8%, mean ± SD, area-label permutation test of the sessions, p = 0.10) and from 10 to 3 percentage points for single-trial decoding (35 ± 3% versus 32 ± 3%, p = 0.03). In Monkey F, FEF and dlPFC did not differ (FEF versus dlPFC, 45 ± 16% versus 56 ± 15% for block decoding and 26 ± 5% versus 31 ± 7% for single-trial decoding, mean ± SD, p = 0.39 and 0.27).

These results were based on activity recorded during successful trials. To test whether the neural representation of task context was related to behavioral performance, we decoded task identity from error trials using the decoder trained on successful trials. We included error trials in which the monkey completed the fixation period but then failed to complete the trial or responded incorrectly (trials in which fixation was broken during the fixation period have no fixation-period activity and could not be analyzed). Error trials were decoded at or only slightly above chance (Figure 3E–F, block-label permutation test of the task within each session). In Monkey H, accuracy with the first four components was 18 ± 3% in FEF and 17 ± 2% in dlPFC (mean ± SD), above chance in FEF with 2, 4 and 5 components (p = 0.045, 0.049 and 0.009) and in dlPFC with 5 components (p = 0.049). In Monkey F, it was 22 ± 4% in FEF (mean ± SD), above chance for every number of components (p = 0.004, 0.003, 0.0004, 0.001 and 0.0005 for 1 to 5 components), and 19 ± 3% in dlPFC, above chance with 1 component only (p = 0.03). Decoding differed between two kinds of error (Figure 3G, first four components). Error trials in which the monkey broke fixation after the fixation period, before the go cue, were decoded below or at chance (12 ± 5% in FEF and 12 ± 6% in dlPFC of Monkey H and 15 ± 2% in FEF and 8 ± 7% in dlPFC of Monkey F, mean ± SD, block-label permutation test of the task within each session, p < 0.0001 in Monkey H, p = 0.43 in FEF and p = 0.0002 in dlPFC of Monkey F), whereas error trials in which the monkey responded incorrectly after the go cue were decoded above chance (22 ± 5% and 21 ± 5% in Monkey H and 28 ± 5% and 28 ± 6% in Monkey F, p < 0.0001 in every area). Response errors were decoded more accurately than fixation breaks in every area (difference in accuracy between the two kinds of error, trial-label permutation test of the error type within each session, sessions combined, p < 0.0001 in both areas of Monkey H and in dlPFC of Monkey F and p = 0.0002 in FEF of Monkey F), but still far less accurately than successful trials. These results suggest that a weaker task signal makes errors more likely, particularly errors made before the go cue.

We next asked how far the population state of each held-out trial moved toward the state of its own task (Supplementary Figure 3A). For every trial, we measured its position along the axis from the mean of all six tasks to the centroid of its own task in the subspace of the first four task components, so that 0 marks the mean of all tasks, that is, no task-specific state, and 1 marks the centroid of the trial’s own task (Supplementary Figure 3B). Held-out successful trials reached 72% of the way to their own task’s centroid in Monkey H and 53% in Monkey F, whereas error trials, on average, did not leave the mean of all tasks (−4% in Monkey H and 0% in Monkey F). The difference between successful and error trials was significant in both monkeys (trial-label permutation test of success and error within each session, p < 0.0001). Interestingly, the error trials were displaced in a systematic direction: error trials of the five spatial tasks were shifted toward the color-task state and error trials of the color task toward the spatial-task states (Supplementary Figure 3C) such that a disproportionate share of the error trials of every task was decoded as the color task (Figure 3F). Furthermore, the position of a single trial on its own-task axis predicted its outcome. The probability of an error decreased as a trial moved from the mean of all six tasks toward the centroid of its own task (Figure 3H, logistic regression with one intercept per session). At the centroid of the trial’s own task, the odds of an error (the error probability divided by the success probability) were 59% lower in FEF and 36% lower in dlPFC of Monkey H and 17% lower in FEF and 34% lower in dlPFC of Monkey F than at the mean of all tasks (trial-label permutation test of success and error within each session and task, p = 0.001 in every area). Therefore, the task-specific population state was weak or absent on error trials, and its strength on a given trial predicted whether the trial would be successful.

One trivial explanation for the task geometry is that the neural representation reflects the color of the fixation cross rather than task identity per se. At one extreme, if the cue color alone drives the fixation-period state, the task would be decodable on trials 1 to 8 of a block, when the fixation cross had a task-specific color, and only at chance on trials 9 to 16, when the fixation cross was white for all six tasks. At the other extreme, if the cue color has no effect on the fixation-period state, then the fixation state should reflect the task performed in the previous trial. To quantify this, we decoded the successful trials of held-out blocks by their position in the block (Figure 3I, first four components). One decoder was trained on the 3rd to 16th successful trials of each training block, so that the decoder was not fit to the first two trials after a task switch. At each position in the block, we measured how often a trial was decoded as its own task and how often as the task of the previous block. As a control, we also measured how often it was decoded as the task of the next block, which the animal could not predict. On the first trial of a block, although the colored fixation cross of the new task was on the screen, activity was decoded as the previous block’s task nearly as often as the new current task (previous versus current task, 29% versus 28% in FEF and 26% versus 24% in dlPFC of Monkey H and 22% versus 23% in FEF and 22% versus 29% in dlPFC of Monkey F, mean across sessions, with SDs of 2–6% across sessions). All 8 values were higher than the control, the rate at which the trial was decoded as the task of the next block (trial-label permutation test, p = 0.0001 in each case in Monkey H and p < 0.04 in Monkey F). These results indicate that the previous task state and the color cue contribute roughly equally to the current task state.

On the second to the eighth trials, when a color cue was presented, the current task was correctly decoded at a relatively constant rate, more often than on the first trial (49 ± 8% versus 28 ± 5% in FEF and 38 ± 9% versus 24 ± 5% in dlPFC of Monkey H and 31 ± 12% versus 23 ± 5% and 39 ± 11% versus 29 ± 6% in Monkey F, mean ± SD across sessions, trial-label permutation test of the trial position within each block, p < 0.0001 in every area). When the fixation cross turned white, decoding of the current task decreased, but far less than the drop to chance expected if the cue alone set the state. In Monkey F, FEF accuracy decreased from 31 ± 12% to 26 ± 7%, and dlPFC accuracy decreased from 39 ± 11% to 32 ± 9% (mean ± SD, trial-label permutation test of the trial position within each block, p < 0.0001 in every area). In Monkey F, the corresponding values were 31 ± 12% to 26 ± 7% and 39 ± 11% to 32 ± 9% (mean ± SD, trial-label permutation test of the trial position within each block, p < 0.0001 in every area). Therefore, both the fixation-cross color and the preceding trials contributed to the fixation-period state, but neither fully explained it: decoding stayed far above chance without the colored cross, and the first trial of a block was already decoded as the new task more often than the control (next block task). The representation thus reflects task context rather than the cue or the previous trials alone.

### Task geometry is conserved across neural populations

Each session sampled a different population of neurons at a distinct recording site (Supplementary Figure 1). We asked whether the task geometry, that is, the arrangement of the 6 task centroids in the task subspace, was shared across neural populations. Because the task subspace of each session is defined by different neurons, only the relative positions of the 6 task centroids, not their coordinates, can be compared between sessions. We first visualized the geometry across sessions by aligning the task centroid configurations (the set of 6 task centroids of a session) of all sessions of an area using generalized Procrustes analysis and averaging the aligned configurations (see Materials and Methods). The resulting common geometry separated the color-mapping, saccade, memory-guided saccade and distractor tasks, whereas the two non-match tasks coincided, in both areas and both monkeys (Figure 4A). Because this alignment rotates every configuration to match the others as closely as possible, it can make the geometries look more alike than they are. To quantify how similar the task geometry was across individual sessions, we compared pairs of sessions within an area with two measures that are invariant to rotation and reflection of the axes. The first was representational similarity analysis (RSA), the Spearman correlation between the pairwise centroid distances of one session and those of another. The second was the Procrustes angle between the two sessions’ centroid configurations, each treated as a single vector of 24 coordinates (6 tasks × 4 components) after both were centered, normalized to unit size and optimally aligned by rotation and reflection. The cosine of this angle, the cosine similarity of the aligned configurations, was used for the statistical tests (see Materials and Methods). In both monkeys, the geometries of two sessions of the same area were more similar than expected by chance (Supplementary Figure 4A and Supplementary Table 1, task-label permutation test of the 6 tasks within every session, p < 0.0001 in Monkey H and p < 0.005 in Monkey F for both measures in both areas). Thus, sessions recorded at different sites shared a common task geometry in both areas of both monkeys.

**Figure 4.**
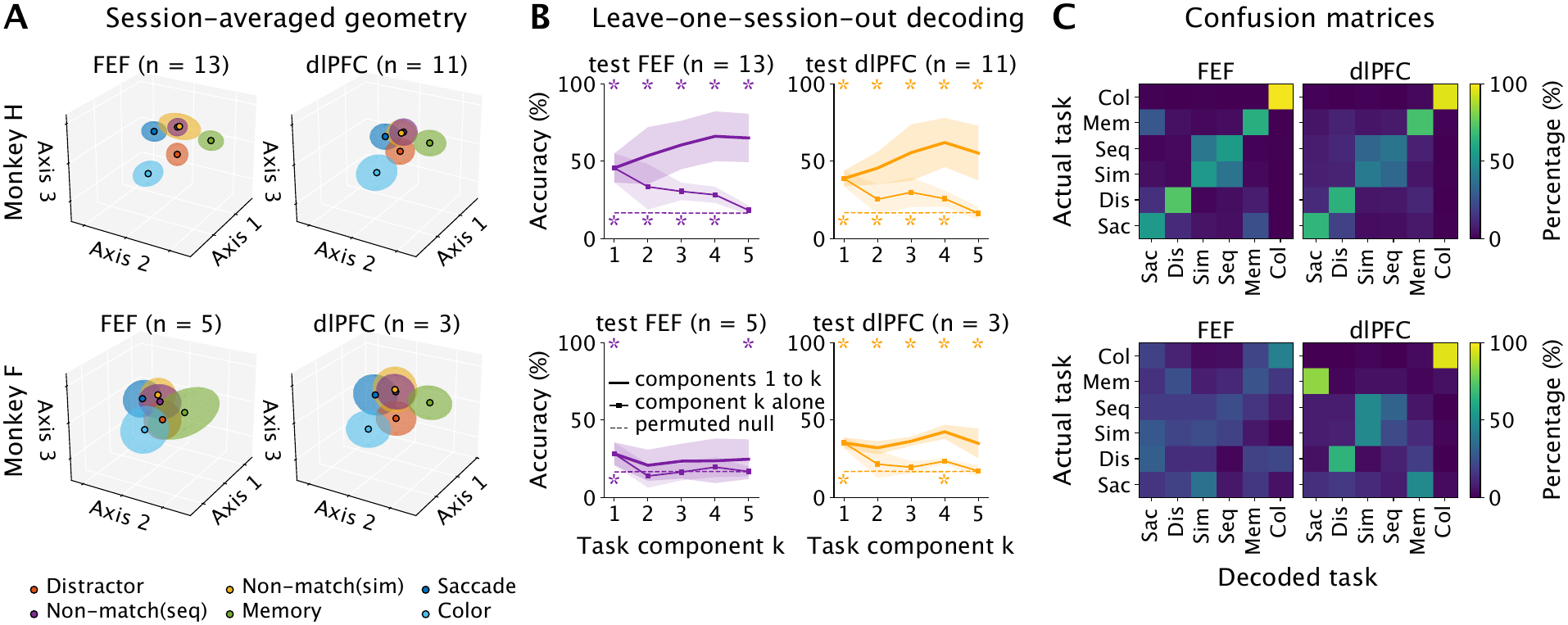
Task geometry is conserved across neural populations. **(A)** Session-averaged task geometry per area and monkey. In each session, every block was projected onto task components estimated without that block (held-out projections), and the 6 task centroids in the first 4 components were aligned across the area’s sessions by generalized Procrustes analysis (see Materials and Methods). For visualization, the first 3 aligned axes are shown. The 5th component, which carried no task information across sessions (B, squares at k = 5), was not used. Dots: centroid of each task, averaged across sessions after alignment. Shading: ±1 SD of the held-out blocks of each task, pooled across the area’s sessions after alignment. n, number of sessions. Monkey H had 13 FEF and 11 dlPFC sessions, and Monkey F 5 FEF and 3 dlPFC sessions. **(B) Leave-one-session-out decoding** of the task blocks of each held-out session from the pooled blocks of all other sessions of the same area, with correspondence-free alignment of the held-out session to the training sessions (see Materials and Methods). Thick lines: decoding from task components 1 to k. Squares: from the k-th axis alone. Lines, squares and shading: mean ± SD across held-out sessions. At each k, the training and test sessions were aligned in components 1 to k, and the squares show the k-th axis of that alignment (see Materials and Methods). Dashed lines: mean accuracy of the null, in which the task labels of every session were permuted independently and the whole analysis was repeated (1,000 permutations, see Materials and Methods). Asterisks: task-label permutation test of the 6 tasks within every session (p < 0.05). **(C) Confusion matrices** of the leave-one-session-out decoding (first 4 components, alignment as in B): session-averaged percentage of held-out blocks of each actual task decoded as each task. Task abbreviations as in Figure 3.

To quantify the consistency of task geometry within an area but across recording sites more stringently, we trained a classifier on the blocks of all but one held-out session within an area, and then asked how well we could decode the blocks of the held-out session (Figure 4B). Each block was represented by its position in the task subspace of its session: the mean activity of its successful trials (z-scored firing rates of all units) projected onto the task components estimated without that block, as in within-session decoding. Because the components of each session are defined by different neurons, these positions cannot be compared directly between sessions. To compare task geometry across sessions, we aligned the training sessions to a common frame in two steps. First, for each training session, we used generalized Procrustes analysis to find the transformation (centering, scaling and rotation) that best matched its 6 task centroids to the consensus, the mean of the aligned task centroids over all training sessions. Second, we applied that transformation to every block of the session and pooled the transformed blocks of all training sessions (the same number of blocks for every task, 23.8 ± 2.2 blocks per task per session in Monkey H and 21.5 ± 2.8 in Monkey F, mean ± SD). We then aligned the held-out session to this frame using half of its blocks and decoded the task of each remaining block from its position in the aligned frame. The two halves then exchanged roles, so that every block was decoded once. Critically, the task labels of the held-out session were used only to group its blocks by task, not to decide which group was which task. Of the 720 possible correspondences of task centroids between training and held-out sessions, we kept the one with the best Procrustes fit, so that in the absence of a shared geometry the expected accuracy is chance (16.7%, see Materials and Methods). In Monkey H, decoding was well above chance in both areas (Figure 4B and Supplementary Table 2, first 4 components, 66 ± 16% in FEF and 62 ± 16% in dlPFC, mean ± SD across held-out sessions, versus 17% for the null obtained by permuting the task labels of every session independently and repeating the whole analysis, task-label permutation test of the 6 tasks within every session, p < 0.001 in both areas), and in 22 of 24 sessions individually. Accuracy across sessions was lower than within sessions (Figure 3A). This could reflect parts of the geometry that are not shared across sessions, since a decoder trained on other sessions can use only the shared part, or an imperfect alignment of the held-out session. Decoding accuracy increased as components were added up to the 4th (gain from 1 to 4 components compared with the gain in the null, p < 0.001 in both areas), and each of the first 4 axes of the common frame decoded the task above chance on its own (Figure 4B, squares, see Materials and Methods). The 5th component neither increased the accuracy further nor decoded the task above chance on its own (Figure 4B, k = 5, and Supplementary Table 2). This is consistent with within-session decoding, which also did not improve beyond the 4th component (Figure 3A and C), because a component that carries no task information within a session has nothing to share with other sessions. As within sessions, the most common confusion across sessions was between the two non-match tasks (Figure 4C). In Monkey F, the task was also decoded across sessions in dlPFC (42 ± 5% versus 17% for the null, p = 0.003) but not in FEF (24 ± 14%, p = 0.055), accuracy did not increase significantly from the first component alone to the first 4 components, and the 5th component again added nothing (Supplementary Table 2). The lower across-session accuracy in FEF of Monkey F than in Monkey H accompanied a weaker task signal within sessions (Figure 3A). FEF test sessions of Monkey F were nevertheless decoded above chance when the training sessions came from dlPFC (Figure 5A).

**Figure 5.**
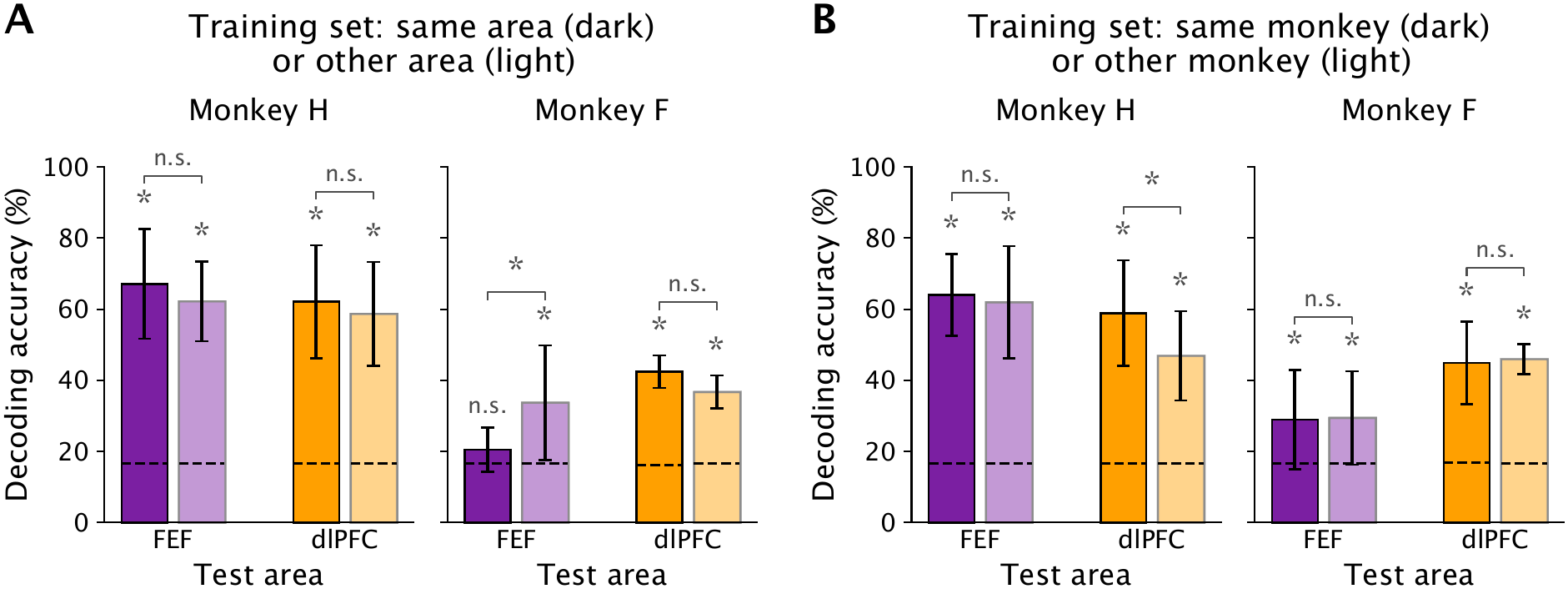
Task geometry is shared between FEF and dlPFC and between monkeys(A) Decoding of each block of the held-out sessions of each test area (FEF on the left and dlPFC on the right) by a decoder trained on other sessions of the same area (dark) or of the other area (light), with the same number of training sessions in every case (10 in Monkey H and 2 in Monkey F). The blocks of the training sessions were aligned to a common frame and pooled, and each block of the test session was assigned to the task whose mean of the pooled training blocks was nearest. The test session was aligned to the training sessions by a one-to-one matching of its 6 task centroids to theirs (720 possible matchings, see Materials and Methods). Bars: mean ± SD across test sessions. Dashed lines: mean accuracy of each bar under the task-label permutation test of the 6 tasks within every session (1,000 permutations). Asterisks above the bars: task-label permutation test of the 6 tasks within every session, p < 0.05 (n.s., not significant). Brackets: same versus other area, area-label permutation test of the sessions. **(B)** Decoding of each block of the same test sessions by a decoder trained on sessions of the same monkey (dark) or of the other monkey (light), with 7 training sessions in every case, aligned and decoded as in A. The training sessions of a Monkey H test session were subsets of the 8 Monkey F sessions or of the 23 other Monkey H sessions, and those of a Monkey F test session were the 7 other Monkey F sessions or subsets of the 24 Monkey H sessions, with accuracies averaged over subsets. Bars: mean ± SD across test sessions. Dashed lines and asterisks are as in A. Brackets: same versus other monkey, monkey-label permutation test of the training sessions.

### Task geometry is shared between FEF and dlPFC and between monkeys

Beyond the consistency within each area, the session-averaged geometries of FEF and dlPFC looked similar to each other (Figure 4A): in both areas and both monkeys, the color-mapping task stood apart, the two non-match tasks coincided, and the remaining tasks were arranged in the same way. To test this similarity quantitatively, we first compared the geometry of every FEF session with that of every dlPFC session (Supplementary Figure 4B and Supplementary Table 3). The two were far more similar than expected by chance (p < 0.0001 for both measures in both monkeys) and nearly as similar as 2 sessions of the same area (in Monkey H, RSA ρ = 0.83 ± 0.07 across areas, mean ± SD over session pairs, versus 0.88 and 0.82 within FEF and dlPFC).

We then asked directly whether a decoder trained in one area could identify the task in the other. To compare training sets fairly, every combination of training and test area used the same number of training sessions, all of whose blocks were pooled for training (Figure 5A, 10 sessions in Monkey H and 2 in Monkey F), and every block of the test session was decoded. In Monkey H, test sessions of both areas were decoded far above chance whether the training sessions came from the same or the other area (Figure 5A and Supplementary Table 2, task-label permutation test of the 6 tasks within every session, p < 0.001 in all 4 cases), and accuracy did not differ significantly between same-area and other-area training (area-label permutation test of the sessions, p = 0.28 and 0.19 for FEF and dlPFC test sessions). In Monkey F, FEF test sessions were decoded above chance with dlPFC training sessions (p < 0.001) but not with FEF training sessions (p = 0.054), and this difference was significant (p = 0.018). With only 2 training sessions, accuracy depends strongly on the quality of the training sessions. The FEF sessions of Monkey F carried a weaker task signal than its dlPFC sessions already within sessions (Figure 3A and C), so 2 FEF sessions gave a noisier training geometry than 2 dlPFC sessions, and the FEF test sessions were decoded better when aligned to the dlPFC geometry. dlPFC test sessions of Monkey F were decoded above chance with both training areas (p = 0.003 and p < 0.001), and the two did not differ significantly (p = 0.61). The fact that each area decoded the other above chance means that FEF and dlPFC shared task geometry, at least in part, in both animals.

In addition to this shared geometry, we asked whether the geometry of a session also carried a signature of its area, with an area identity index (Supplementary Figure 5, first 4 components). Each session was left out in turn and assigned to FEF or dlPFC according to which area’s mean geometry, computed from the other sessions, better matched the geometry of the left-out session. In Monkey H, 18 of 24 sessions (75%) were assigned to their own area, versus 50% when the area labels were shuffled across sessions (area-label permutation test of the sessions, p = 0.06), and the mean geometries of FEF and dlPFC differed with a similar trend (p = 0.06). In Monkey F, neither measure departed from chance (1 of 8 sessions (13%) assigned to its own area, p = 0.12, and p = 0.10 for the difference between the area means). Because Monkey F contributed only 8 sessions, compared with 24 in Monkey H, this null result may reflect the smaller data set rather than the absence of an area signature. Thus, in addition to the shared geometry, FEF and dlPFC may each carry a smaller area-specific component, which the present sessions were too few to establish.

We next asked whether the geometry was also shared between animals. The task geometries of a Monkey H session and a Monkey F session were far more similar than expected by chance (Supplementary Figure 4C and Supplementary Table 3, RSA ρ = 0.69 ± 0.18, mean ± SD over 192 session pairs, p < 0.0001 for both measures). We then decoded the blocks of each monkey’s sessions with a decoder trained on sessions of the other monkey and compared it with a decoder trained on the same number of sessions of the same monkey (Figure 5B, 7 training sessions for every decoder). Both decoders were above chance in every case (Supplementary Table 2, task-label permutation test of the 6 tasks within every session, p < 0.005), and across-monkey decoding was as accurate as within-monkey decoding (monkey-label permutation test of the sessions, p > 0.5) except in dlPFC of Monkey H, where it was somewhat less accurate (47 ± 13% versus 59 ± 15%, mean ± SD across test sessions, p = 0.027). These results suggest that the task geometry was also shared between monkeys.

### Global signals do not fully account for the task geometry

Finally, we asked what the conserved task geometry reflects. One possibility is that the geometry reflects global signals that differ between tasks but are not specific to the content of any task, such as the expected reward, the difficulty of the task or the level of attention. If so, the positions of the tasks along the task components should follow the amount of reward received for each task, the hit rate (a measure of difficulty) or the saccade reaction time (a measure of attention). We tested this in all 32 sessions of both monkeys, because the geometry was shared across sessions, areas and animals (Figure 6). For each of the first 4 task components, we averaged the distances between the 6 tasks along the component over sessions and correlated them with the differences between the tasks in each variable, also averaged over sessions (Spearman correlation over the 15 task pairs). Because the sessions had similar task geometries, they do not provide independent evidence about a variable. Whether the geometry follows a variable depends on how the variable is distributed over the 6 tasks, not on how many sessions were recorded. We therefore shuffled the 6 task labels in the same way in every session, recomputed the correlation for each of the 720 possible shuffles, and compared the observed correlation with this distribution (see Materials and Methods). Reward amount correlated with the first component (ρ = 0.77, p = 0.017), although not significantly after correction across 12 tests, one for each combination of the 3 variables and 4 components (FDR-corrected p = 0.20), and not with components 2 to 4 (p > 0.1). The color-mapping task was also the most rewarded, so this correlation cannot be distinguished from the separation of the color task from spatial tasks along the first component. Hit rate and saccade reaction time did not correlate with any component after correction (all FDR-corrected p > 0.2). Thus, global signals did not fully account for the task geometry, which may also reflect more task-specific variables, such as the relevant sensory modality, the demand on spatial memory or the planned saccade (see Discussion).

**Figure 6.**
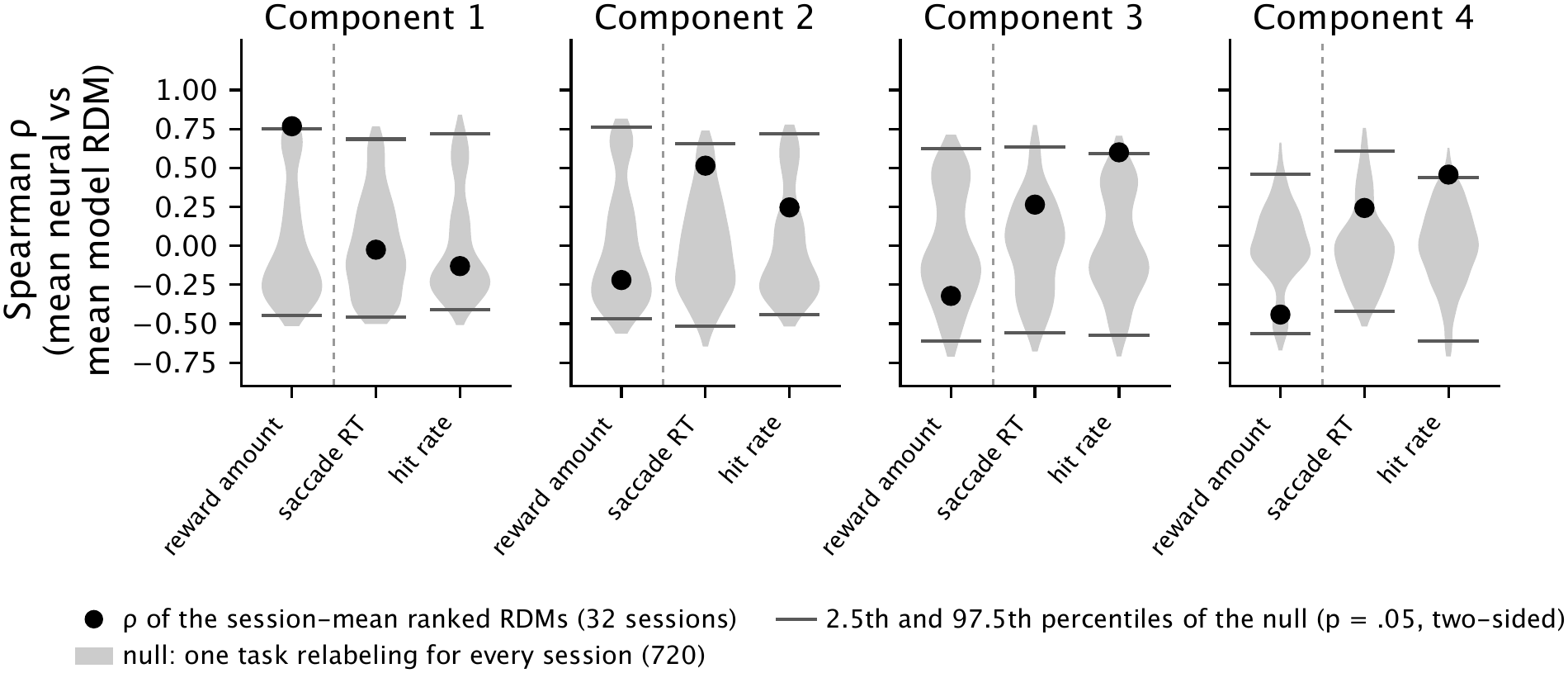
Global signals do not fully account for the task geometry. Correlation between the task geometry along each of the first 4 task components and the differences between the tasks in reward amount (total reward per trial), saccade reaction time (ms), and hit rate (%). For each component, the neural dissimilarity between two tasks was the distance between their centroids along that component of the session, and the model dissimilarity was the absolute difference between the two tasks in the variable. The 15 neural and the 15 model dissimilarities were rank-transformed in each session and averaged over sessions, and the two averages were compared by Spearman correlation. Black dots: Spearman correlation. Gray violins: null distribution of the correlation when one permutation of the 6 task labels is applied to every session (all 720 permutations). Dark ticks: the 2.5th and 97.5th percentiles of the null. No correlation reached p < 0.05 after FDR correction across the 12 tests (3 variables and 4 components). All 32 sessions of both monkeys, 13 FEF and 11 dlPFC sessions in Monkey H and 5 and 3 in Monkey F.

## Discussion

Flexible behavior may be facilitated by configuring task-appropriate “control states” of the brain before the moment-to-moment computations of a trial unfold^5^. In the present work, we leveraged large-scale spiking recordings, with hundreds of neurons recorded simultaneously in FEF or dlPFC, while monkeys performed 6 interleaved tasks to examine whether and how population activity is systematically modulated by task contexts even before a trial begins. We found that frontal cortical neurons adjusted their baseline activity for the upcoming task within a task subspace, in which each component separated different sets of tasks, and that the arrangement of the 6 task states in this subspace was conserved across neural populations from different recording sessions, across the two areas, and across the two animals. Notably, the task representation persisted even when an explicit sensory cue for task identity was removed (i.e., when the task-specific color cue was absent in the second half of trials within each block), implying that the neural population can maintain an internally defined task context. This reconfiguration of baseline population activity may alter how the frontal cortex interacts with any other area, whether upstream, downstream or at the same level of the hierarchy, and may also change the dynamics within each area, so that the same incoming signals are processed differently in different tasks, thereby supporting task-specific adaptive behavior^14^. Such preparatory states need not be confined to the frontal cortex. In mice performing a task in which the probability of the stimulus side switched between blocks, the block prior was encoded in activity before stimulus onset in 20 to 30% of brain regions, from early sensory to motor and high-level cortical areas^15^. Whether the task states found here in FEF and dlPFC extend to other areas, with the same geometry, remains to be tested.

The existence of a shared task geometry is itself notable. When a neural representation is driven by an input that varies continuously along a few dimensions, it is not surprising that the representation spans the same number of dimensions and preserves the structure of the input. Responses to spatial targets in many cortical areas form a map of space in some neural subspace, from which the location of a target can be read out, and representations of color may likewise occupy a subspace whose geometry mirrors color space. Space and color, however, have an inherent structure that a representation can inherit. Tasks do not: they are discrete rules, learned by the animal, with no obvious distance between them. Therefore, it is remarkable that the fixation-period activity of the frontal cortex arranged the 6 tasks in a consistent geometry from which task identity could be read out, and more remarkable still that this geometry was shared by populations recorded in different sessions, in FEF and dlPFC, and in two monkeys (Figures 4 and 5 and Supplementary Figure 4). A shared geometry could arise if tasks have an inherent structure, set by the cognitive operations they share and those that distinguish them, which the frontal cortex represents in a common way across individuals. Global signals such as expected reward, task difficulty, and attention, measured by reward amount, hit rate and saccade reaction time, did not fully account for the geometry (Figure 6), consistent with a structure set at least in part by the content of the tasks rather than by how rewarding, difficult or engaging they were.

Which task-specific variables shape the geometry remains open. The color-mapping task, the only task in which color was relevant, was the most separated from the others, and the two non-match tasks, which share a response rule, coincided, consistent with a geometry organized by the content of the tasks, such as the relevant sensory modality, the demand on spatial memory or the planned saccade. With 6 tasks, however, such variables cannot be tested statistically. Most of them take only two or three values across the tasks: the relevant sensory modality, whether the saccade target was visible at the go signal, and the number of initial targets each single out one task, spatial memory demand separates three tasks from the other three, the central color cue and the number of total targets separate two tasks from the other four, and the number of choice options takes three values. Therefore, the values of those variables can be assigned to the tasks in only 6 to 60 distinct ways, so no comparison across these tasks can reach significance after correction, and the color-mapping task differed from the others in reward amount and trial length as well as in sensory modality. Designs with more tasks, in which such variables vary independently of one another, will be needed to identify them.

Although we observed a consistent neural geometry for the 6 tasks across time and neural populations, an important open question remains: whether this geometry is shaped by the particular set of tasks the animals have learned and performed together. For example, would adding a seventh task change the neural geometry for the original 6 tasks? Similarly, would reducing the set to 5 tasks change the population activity and its neural geometry for those remaining tasks? Addressing these questions by experimentally manipulating the composition of the task set would be an important direction for future work. Such experiments would reveal whether task representation is inherently relative (shaped by the full repertoire of tasks the animal has internalized).

While our findings suggest a link between preparatory neural states and task performance, the nature of this relationship remains complex. We found that task information was strongest in trials in which animals maintained fixation to the end of the trial and then made a correct response, present but weak when fixation was maintained but an incorrect response was made, and not present in trials when animals held fixation at least until the first stimulus appeared but then broke fixation before the end of the trial (Figure 3G). This is consistent with an appropriate preparatory state contributing to successful performance^4^. However, other interpretations are possible. Behavioral errors can arise through multiple routes. In addition to a failure to set the correct task set, a lapse in attention or miscoding of stimulus information could also lead to an error. Moreover, even if correct task representation in the frontal cortex is necessary for successful performance, it need not occur during the preparatory period and could emerge later, during a delay period or at the moment a response is required. Thus, preparatory neural states may be one of several contributing factors to successful task performance, rather than a strict requirement.

## Materials and Methods

Two male rhesus monkeys (Macaca mulatta), F and H, were used in this study. All procedures complied with the Guide for the Care and Use of Laboratory Animals and were approved by the Washington University Institutional Animal Care and Use Committee.

During experimental sessions, the animals were seated in a custom primate chair (Crist Instrument, MD, USA) with the head immobilized via an implanted head post (Graymatter Research, MT, USA) in a completely darkened room. Visual stimuli were generated with custom software and displayed using an organic light-emitting diode television (LG, South Korea). Eye position was recorded with an infrared video eye-tracking system (ISCAN, MA, USA).

### Behavioral paradigms for six tasks

Monkeys were trained to perform six visual saccade tasks that required making a saccadic eye movement to a task-specific target on the screen. In every task, each trial began with a 1-second fixation period during which the animals were required to maintain gaze on a central fixation cross on the screen. During this period, only the fixation cross appeared, and its color indicated the current task (but see below). When the peripheral targets appeared, they were the same color as the fixation cross (except in the color-mapping task and for the early task-irrelevant color cue in the saccade task). The animals were required to maintain gaze on the central fixation cross until it disappeared. A liquid reward was delivered when the gaze reached near the correct location on the screen within 150 milliseconds after leaving the fixation cross. Every trial in which the reward was not delivered was counted as an error trial, whether the animal broke fixation during the fixation period or failed later in the trial.

Tasks were organized in blocks of 16 successful trials. After 16 successful trials, the task switched randomly to another task, including the possibility of repeating the same task. Within each block, the fixation cross and peripheral stimuli were presented in the task-specific color during the first 8 trials and in white during the last 8 trials in all but the saccade and color-mapping tasks. In the saccade task, the fixation cross and the peripheral saccade target were white during the last 8 trials of each block. In the color-mapping task, the fixation cross was white.

See below for a detailed description of each task.

(1) ***Distractor task***: Following the 1-second fixation period, a peripheral target appeared for 200 milliseconds at a random location on an invisible circle with a radius of 13 degrees of visual angle (dva) centered on the fixation cross and then disappeared. After a delay period of 1.01– 1.51 seconds, during which only the central fixation cross remained on the screen, a second peripheral target appeared for 200 milliseconds and was followed by a second delay period of 0.51–1.51 seconds. Finally, a third peripheral target appeared and then the fixation cross disappeared. The animals were required to make a saccade to the last peripheral target to receive a liquid reward. The first target could appear anywhere on the circle. The third target was separated from the first two targets by ±60°, ±120°, or 180°. Thus, there were four relative configurations of the three targets, but over 1,440 absolute configurations. During the first 8 trials of each block, the fixation cross and peripheral targets were colored grey.
(2) ***Non-match-to-sample with sequential two-item task***: Identical to the distractor task until the end of the second delay period, except that the fixation cross and peripheral targets were red during the first 8 trials of each block. After the second delay period, the two previously presented targets re-appeared on the screen along with a third novel target. When the fixation cross disappeared, the animals were required to make a saccade to the novel target.
(3) ***Non-match-to-sample with simultaneous two-item task***: Similar to the sequential two-item non-match-to-sample task, except that the first two targets were presented simultaneously, followed by a single longer delay period (2.03–2.53 seconds), such that total trial time was roughly matched. As in the sequential non-match task, the fixation cross and peripheral targets were red during the first 8 trials of each block. Because of this, the first trial of a new block was not dissociable between the two non-match tasks during the fixation period. However, based on the block structure, the remaining 15 trials were predictable as belonging to one of the two tasks.
(4) ***Memory-guided saccade task***: Following the fixation period, a peripheral target appeared for 200 milliseconds at a random location on an invisible circle with a radius of 13 dva centered on the fixation cross and then disappeared. After a delay period of 2.1–2.6 seconds, the fixation cross disappeared, and the animals were required to make a saccade to the previously presented target location while no visual stimuli were present on the screen. The task-specific color for the fixation cross and peripheral targets was green.
(5) ***Saccade with a task-irrelevant color cue task***: After the fixation period, a square colored either cyan, blue, or yellow appeared behind the fixation cross at the center of the screen. After a delay of 0.72–1.22 seconds (cue period), a peripheral target appeared at a random location on an invisible circle with a radius of 10 dva, and the fixation cross disappeared. The animals were required to make a saccade to the peripheral target while it was present. Brown was used as the task-specific color.
(6) ***Color-mapping task***: Identical to the saccade task until the end of the cue period. At this time, two colored targets appeared 10 dva from the fixation cross, above and to the left and right of it: one in the color paired with the central color cue and the other in one of the two remaining colors (cyan, blue, or yellow). The animals were pre-trained on arbitrary color–pair associations (blue→cyan, cyan→yellow and yellow→blue for Monkey F, and blue→blue, cyan→cyan and yellow→yellow for Monkey H). Based on these associations, the animals were required to make a saccade to the peripheral target whose color was paired with the central color cue, regardless of its spatial location. Which target appeared on the left and which on the right was randomly assigned on each trial.

### Electrophysiology

In each experimental session, we lowered a Neuropixels 1.0 NHP probe into the frontal eye field (FEF) or the dorsolateral prefrontal cortex (dlPFC) (Supplementary Figure 1). After allowing a minimum of 45 minutes for stabilization, we recorded spiking activity using the Open Ephys system while the monkeys performed the six tasks. In 9 of the sessions of Monkey H, a seventh task (a pop-out visual search task) was interleaved with the six tasks in the same block design, but its trials were not analyzed. Units were isolated with Kilosort 4^16^ and curated by considering categories from Bombcell^17^. To exclude clusters driven by non-neuronal signals, we included only units that responded to the presentation of a stimulus. We also required a mean firing rate of at least 0.5 spikes/s during the fixation period. The resulting dataset comprised 8,934 units from 24 sessions in Monkey H (13 FEF sessions with 338–618 units each and 11 dlPFC sessions with 112–416 units) and 2,559 units from 8 sessions in Monkey F (5 FEF sessions with 106–543 units and 3 dlPFC sessions with 134–486 units).

All analyses used the fixation period only: for each unit and trial, we took the mean firing rate over the last 800 ms of the fixation period, dropping the first 200 ms to avoid transients associated with the start of fixation. (The fixation period of the color-mapping task of Monkey H was shorter, 700 ms rather than 1000 ms, and so only the last 500 ms were used in that case.) For most analyses we included only successful trials. For the error analysis, we also included error trials in which the error occurred after the fixation period (10,724 of 18,022 error trials in Monkey H, 60%, 368–575 per session, and 2,066 of 4,086 in Monkey F, 51%, 199–291 per session). Of these, only the error trials of the blocks kept for decoding (see below) entered the neural analyses (10,415 in Monkey H, 295–575 per session, and 2,031 in Monkey F, 193–284 per session). Error trials in which fixation was broken during the fixation period are counted as errors in the hit rates (Figure 1) and in the trial numbers from the block transition (Figure 1H) but were excluded from all neural analyses.

### Task components and task geometry

Task components were estimated with regularized linear discriminant analysis (LDA). For each session, the firing rates of successful trials were averaged within each block (one population vector per block) and z-scored per unit, and the five discriminant axes that maximize the between-task relative to the within-task (across-block) variance were obtained by solving the generalized eigenvalue problem with the pooled within-task covariance regularized by Ledoit– Wolf shrinkage^18^. To estimate the task components, we used block-averaged activity because the task context was set at the level of blocks: variability across blocks of the same task reflects the variability of the task state, whereas trial-to-trial variability within a block is due to factors other than the task that is about to be performed, since that is held constant. The axes were orthonormalized in the order of their eigenvalues, so that projections are in units of z-scored firing rate and are referred to as task components 1–5. Each subspace spanned by the first k components is a “task subspace”. (With six tasks, LDA can extract at most five discriminant dimensions.) The task geometry of a session was defined as the configuration of its six task centroids in the task subspace (Figure 2B). Unless stated otherwise, summary statistics use the first four components, the number beyond which decoding accuracy no longer increased within or across sessions (Figures 3A, 3C and 4B). The task geometry of single sessions (Figure 2B) and the session-averaged geometry of Figure 4A were built from the held-out projections, the same as used for decoding. Because the sign and orientation of the components are arbitrary, all panels were rotated, with reflection allowed, into the frame of one reference geometry. The reference was the generalized Procrustes consensus of all 32 sessions of both monkeys, built after the centroid configuration of every session was centered and normalized to unit size.

Generalized Procrustes analysis determines the shape of the consensus, and the consensus keeps the orientation and component order of the session where the alignment starts. The rotation of each panel was found after its configuration was normalized in the same way. In Figure 2B, this rotation was then applied to the unnormalized projections, so that the axes keep units of z-scored firing rate. In Figure 4A, it was applied to the session-averaged geometry of each area of each monkey. To obtain this geometry, the sessions of the area were aligned by generalized Procrustes analysis, which brings every session to the same size, and then averaged. The rotation can mix the components and not only flip their signs, but it does not change the distances between task centroids. The reference sets only the orientation of the figures and was not used in any quantitative analysis. All 3D panels share one viewing angle. Shading shows the SD of the single trials of each task around their centroid in Figure 2B (trial-to-trial variability within the session, with each held-out trial projected onto the components estimated without its block) and, in Figure 4A, the SD of the held-out blocks of each task, pooled across the area’s sessions after each session’s blocks were mapped with that session’s alignment.

### Within-session (within-population) task decoding

Within-session decoding used leave-one-block-per-task-out cross-validation. In each repetition, one block of every task was held out as the test set (Figure 3). Whole blocks rather than single trials were held out because slow drifts made trials of the same block more similar to each other than to trials of other blocks of the same task (Supplementary Figure 2). In each session, every task kept the same number of blocks, the smallest number of blocks of any task (16 to 29 across sessions), and the extra blocks of the other tasks were dropped at random and not used for decoding or task geometry. The number of repetitions equaled this number of blocks, so that every training set lacked exactly one block of each task and contained the same number of blocks of every task. Blocks were selected for testing in random order within each task. Five task components were estimated with LDA from the block means of the remaining blocks (the training set), and, in different analyses, either the held-out block means, the held-out successful trials, or the error trials of the held-out blocks were projected onto the first k components (k = 1 to 5) and assigned to the task with the nearest training centroid (Euclidean distance). Training centroids were computed from block means for block decoding and from single trials for single-trial decoding. To compare FEF and dlPFC with equal numbers of neurons, every session was also decoded with 100 randomly drawn neurons (10 draws, accuracies averaged over draws), and the areas were compared by permuting the area labels of the sessions (10,000 random relabelings in Monkey H and all C(8, 3) = 56 relabelings in Monkey F). For decoding by position in the block (Figure 3I), all blocks of a session with at least 8 successful trials were divided in chronological order into 8 folds of consecutive blocks, and the 2 blocks on either side of the held-out blocks were left out of training, because slow drifts made neighboring blocks similar and the block preceding a held-out block would otherwise be in the training set. Each training set kept the same number of blocks of every task, with the extra blocks dropped at random. Task components and centroids were estimated from trials 3 to 16 of the training blocks, which include both cue conditions, and every successful trial of the held-out blocks was decoded with the first four components. At each position in the block, we computed the percentage of trials decoded as their own task, as the task of the previous block and as the task of the next block. The previous-and next-block readouts used only blocks that followed or preceded a switch to a different task, excluding switches between the two non-match tasks, which share the red fixation cross. Because each trial was assigned to exactly one of the six tasks, and the previous and next tasks always differed from the current task, a trial decoded as its own task could not also be decoded as the previous or next task. Chance for the previous-and next-block readouts was therefore not 16.7% but the trials not decoded as their own task divided among the five other tasks, (1 − own-task accuracy)/5, about 12%, and it varied with own-task accuracy across positions in the block. For this reason, the next-block readout, which the animal could not predict, rather than a fixed chance level, served as the baseline for the previous-block readout. On the first trial, the previous-block readout and the own-task readout were each compared with the next-block readout by swapping the two task labels at random within each block (1,000 permutations per session). Own-task decoding on trials 2 to 8 versus trial 1 and versus trials 9 to 16 was compared by permuting the trial positions within each block (1,000 permutations per session). Sessions were combined as described under Statistics. For the null distribution, task labels were permuted across blocks (100 permutations per session) and the whole procedure was repeated. Whether adding a component increased decoding was tested with a one-sided test for each increment from k − 1 to k components (Figure 3A and C, brackets), because the question was whether the added component carried task information. The statistic was the mean over sessions of the gain in accuracy, and its null distribution was the gain computed within each block-label permutation, with one permutation drawn independently for every session (10,000 draws).

To show the slow drift and its effect on decoding (Supplementary Figure 2), we used the blocks kept for decoding. The successful trials of every block were split at random into two halves of equal size, and the Euclidean distance between the mean z-scored firing rates (all units) of the two halves of a block was compared with the distance between a half of that block and a half of another block of the same task (Supplementary Figure 2A). Drift was measured in each session as the Spearman correlation between the distance of the mean activity of two blocks of the same task and the time between them, the difference between their mean trial numbers in the session (Supplementary Figure 2B). It was tested against a null in which the time positions of the blocks were shuffled within the session (1,000 shuffles per session), with sessions combined as described under Statistics. To show the effect of the cross-validation design (Supplementary Figure 2C), single-trial decoding with whole blocks held out, as above, was compared with decoding in which single trials were held out at random within each task regardless of their block, so that the other trials of their blocks stayed in the training set. Both designs used the same number of folds, the first four components, and the block-label permutation null (50 permutations per session). The paired differences between the two distances and between the two designs were tested across the sessions of each monkey with a two-sided sign-flip test, which used all 256 sign assignments in Monkey F and 10,000 random assignments in Monkey H. FEF and dlPFC were pooled because the 3 dlPFC sessions of Monkey F alone allow no p-value below 0.25.

Task signal strength (Figure 3H and Supplementary Figure 3B) is the projection of a held-out trial onto the direction from the origin of the task subspace (first four components from training set of successful trials) to the centroid of its own task, expressed as a fraction of the centroid’s distance (0 for no signal toward its own task and 1 for as far as the centroid). The mean task signal strength of successful and error trials (Supplementary Figure 3B) was compared by shuffling the success and error labels of the held-out trials within each session (1,000 permutations) and combining sessions as described under Statistics. The relationship between task signal strength and trial outcome (Figure 3H) was quantified by logistic regression of the outcome of every held-out trial (error or success) on its task signal strength, pooling the trials of all sessions of an area with a separate intercept for each session. The slope was tested against a null distribution obtained by shuffling the success and error labels of trials within each session and task (1,000 permutations). For Figure 3G, the held-out error trials with the complete fixation window were divided by the kind of error, classified from the eye position: fixation breaks after the first task stimulus and before the go cue (990 trials in Monkey H, 41 ± 11 per session, mean ± SD, and 432 trials in Monkey F, 54 ± 18 per session) and response errors after the go cue, that is, saccades that landed nearest a wrong target or missed the correct target (4,831 trials in Monkey H, 201 ± 55 per session, and 1,055 trials in Monkey F, 132 ± 44 per session). The other error trials (no response after the go cue, fixation breaks caused by blinks, and unclassified trials) were not included. Decoding accuracy of each kind was computed in every session with at least 5 trials of that kind and compared with the null of the block-label permutation test of the task within each session, computed for the error trials, and the two kinds were compared by shuffling the trials between the kinds within each session (1,000 permutations), combining sessions as described under Statistics. For Supplementary Figure 3C, each held-out trial was projected onto the direction from the mean of the successful spatial-task trials (0) to the centroid of the successful color-task trials (1), and the difference between successful and error trials was tested by permuting the trial labels within each session (1,000 permutations) and combining sessions as described under Statistics.

### Across-session (across-population) task geometry comparisons

Task geometries were compared between sessions in the subspace of the first four components. Representational similarity^19^ was the Spearman correlation between the 15 pairwise Euclidean distances among the six task centroids of one session and those of another. Cosine similarity was the cosine of the Procrustes angular distance^20^. The six task centroids of a session in the first four components were treated as one configuration, a 6 × 4 matrix X with one row per task, so that the whole configuration of one session was compared with that of another rather than one centroid at a time. Each configuration was centered by subtracting the mean over the six tasks and scaled to unit Frobenius norm (ǁXǁ = 1). The Procrustes distance between two sessions X and Y is the smallest value of ǁX − YRǁ over all orthogonal matrices R, that is, over all rotations with reflection allowed, where ǁX − YRǁ is the square root of the summed squared distances between the centroids of the same task in the two configurations. The minimizing R was obtained from the singular value decomposition of YᵀX. Treated as vectors of 24 coordinates (6 tasks × 4 components), X and YR both have length 1, because both configurations were scaled to unit norm and rotation does not change length. The squared Procrustes distance therefore depends only on the angle θ between the two vectors, ǁX − YRǁ² = 1 + 1 − 2 cos θ = 2 − 2 cos θ, so the R that minimizes the Procrustes distance is the R that maximizes cos θ. This maximal cosine, cos θ = tr(XᵀYR), which is the sum of the products of the matching coordinates of X and YR, is the cosine similarity, and θ is reported as the Procrustes angle (Supplementary Figure 4). For each pair of sessions, the null distribution was obtained by randomly reassigning the six task labels among the tasks of each session (the same reassignment for all blocks of a session, independently for the two sessions) and recomputing the measure. Because both measures depend only on how one session is relabeled relative to the other, this distribution was computed exactly from all 720 reassignments of the task labels of one session. For the group-level test (Supplementary Figure 4A), the task labels of every session of an area were permuted independently and the mean similarity over all pairs was recomputed (10,000 permutations). The same test was applied to the pairs of an FEF and a dlPFC session (Supplementary Figure 4B) and to the pairs of a Monkey H and a Monkey F session (Supplementary Figure 4C). For the session-averaged geometry (Figure 4A), the centroid configurations of all sessions of an area were aligned by generalized Procrustes analysis^21^, which aligns every session to the mean of all sessions rather than to any single session, so that the shape of the averaged geometry does not depend on which session is chosen as the reference, only its orientation. The configuration of the first session of the area served only as a starting reference, and each iteration had two steps. First, every configuration was rotated, with reflection allowed, to minimize its Procrustes distance to the reference. Second, the mean of the rotated configurations, rescaled to unit norm, became the new reference. The two steps were repeated until the reference no longer changed, and the aligned configurations were then averaged to give the session-averaged geometry. The same procedure aligned the training sessions in the cross-session decoding.

For cross-session task-block decoding (Figures 4B–C, 5A and 5B), each session in turn served as the test session and was decoded by a decoder trained on other sessions. Each block was represented by one point: the mean z-scored activity of its successful trials, projected onto the task components of its own session (estimated without that block, as in within-session block decoding). Points from different sessions cannot be pooled directly, because each session has its own neurons and its components have an arbitrary orientation. The training sessions were therefore first put into a common frame, in two steps. First, the six task centroids of each training session (one per task label) were aligned across sessions by generalized Procrustes analysis. The consensus was the mean of the aligned centroids of all training sessions. The consensus was found iteratively: every session was aligned to the current mean, the mean was recomputed, and the two steps were repeated until the mean no longer changed. This gave one transformation per session: centering, scaling to unit norm and rotation, with reflection allowed. Second, each session’s transformation was applied to all of its block points, and the transformed blocks of all training sessions were pooled to form the training set. The test session’s blocks were split into two halves, and the six centroids of one half were matched to the six tasks of the training consensus in every possible way (720 matchings). For each matching, the centroids were centered, normalized to unit size and rotated, with reflection allowed, to best match the consensus, and the matching with the highest cosine similarity was kept. Critically, the test session’s task labels were thus used only to group its blocks into six clusters and to score the decoding, never to decide which cluster was which task. The blocks of the other half were then mapped into the common frame with this same transformation, and each block was assigned to the nearest training centroid (the mean of the pooled training blocks of a task). The two halves then exchanged roles, so that every block of the test session was decoded once. For Figure 4B, the alignment was redone for each number of components k, using only components 1 to k of every session, so that the training and test sessions shared one frame of k axes. The test blocks were then decoded either from all k axes or from the k-th axis alone. The orientation of this frame is set by the first training session, from which the alignment starts, so its axes resemble that session’s components but are not identical to them, because the reference is updated to the mean of all sessions as the alignment proceeds. Critically, decoding from k axes does not depend on this orientation, because rotating the common frame leaves the distances between blocks and task centroids unchanged. All cross-session analyses, other than Figure 4B, used the alignment in the first 4 components, which is the alignment at k = 4 in Figure 4B. For the null distributions of Figures 4B, 5A and 5B, the task labels of every session (in Figure 5, the sessions of both monkeys) were permuted independently (the six tasks were reassigned among themselves, so that all blocks of a task kept one shared label), and every step was repeated, from the generalized Procrustes alignment of the training sessions to the choice of correspondence and the decoding (1,000 permutations). The group-level p-value was the fraction of permutations in which the mean accuracy over the test sessions was at least as far from the average of all permutation means as the observed mean accuracy, in either direction. The changes from 1 to 4 and from 4 to 5 components in Figure 4B were tested in the same way, with the mean change over the test sessions as the statistic. To compare training areas without a difference in training-set size (Figure 5A), each test session was decoded from the same number of training sessions whether they came from its own area or the other area. This number was set by the smaller area, dlPFC (10 training sessions in Monkey H and 2 in Monkey F). When more sessions than this were available, the training set was subsampled: all possible subsets were used if there were at most ten, otherwise 10 randomly chosen subsets, and the accuracies were averaged over subsets. For the comparison across monkeys (Figure 5B), same-monkey and other-monkey decoders were trained on 7 sessions drawn regardless of area, the number of sessions left in Monkey F when one of its sessions is tested. When more than 7 sessions were available, training subsets were drawn and their accuracies averaged.

The area identity index (Supplementary Figure 5) asked whether the task geometry of a session revealed the area in which the session was recorded. The geometry of each session was summarized by the 15 distances between its 6 task centroids, computed from the held-out block projections in the first four components. To keep noise from inflating the distances, the blocks of each task were split at random into two halves, and each squared distance was computed across the two halves, as the product of the centroid difference in one half and the same difference in the other half, averaged over 200 random splits. The 15 distances were then divided by their mean, so that only the shape of the geometry was compared. No alignment between sessions was needed, because distances do not depend on the orientation of the components. Each session was left out in turn and assigned to FEF or dlPFC, whichever area’s mean geometry, computed from the other sessions, better matched the geometry of the left-out session. The match was measured with a Gaussian likelihood, using the within-area covariance pooled over the two areas and shrunk with the Ledoit–Wolf method, and equal prior probabilities for the two areas. The statistic was the fraction of sessions assigned to their own area. For its null distribution, the area labels were shuffled across sessions, keeping the number of sessions in each area, and the whole procedure was repeated (1,000 shuffles). As a second test, the Mahalanobis distance between the FEF and dlPFC mean geometries, with the same covariance, was compared with its distribution over 10,000 shuffles of the area labels.

### Correlations with reward amount, reaction time and hit rate

To ask whether the task geometry reflected global signals (Figure 6), for each session and each of the first 4 task components, we compared the distances between the six tasks along the component with the differences between the tasks in three variables measured in that session: reward amount per trial, saccade reaction time, and hit rate. The neural dissimilarity matrix contained the absolute differences between the coordinates of the six task centroids on the session’s own k-th component (15 task pairs). This requires no alignment across sessions because a one-dimensional dissimilarity matrix does not depend on the sign or the scale of the axis. The model dissimilarity matrix contained the absolute differences between the tasks in the variable.

As in representational similarity analysis^19^, the neural and model dissimilarity matrices of each session were rank-transformed and averaged over all 32 sessions of both monkeys, and the two mean matrices were compared by Spearman correlation. Each variable was tested on its own rather than in a multiple regression, because 15 task pairs that derive from only 6 tasks are too few to separate the contributions of several variables. Group-level significance was assessed with a common task-label permutation test: the same permutation of the six task labels was applied to the neural geometry of every session, which is equivalent to permuting the rows and columns of the mean neural dissimilarity matrix together, and the observed correlation was compared with its distribution over all 720 permutations. Because the sessions share one task geometry, permuting the labels of each session independently would treat the sessions as independent evidence for a variable, whereas the evidence rests on the six tasks. The test was two-sided and p-values were corrected across the 12 tests (3 variables × 4 components) with the Benjamini– Hochberg false-discovery-rate (FDR) procedure. The smallest possible p-value of this test is 1 over the number of distinct ways in which the values of a variable can be assigned to the six tasks. Reward amount, saccade reaction time, and hit rate differed among all six tasks, so their smallest possible p-value was 1/720 ≈ 0.0014 before multiple comparison correction.

## Statistics

Statistical tests were permutation tests, except for the Friedman test of firing rates (Figure 2A), the one-way ANOVA of block-mean rates used to identify single units whose fixation-period firing rate differed among tasks (p < 0.01), and the paired t-test used to select visually responsive units. The null hypothesis of each permutation test determined which label was permuted. Block-label permutation tests permuted the task labels across the blocks of a session (within-session decoding). Trial-label permutation tests permuted a label of the trials, namely success/error or error type within a session, and cue stage, trial position, or the previous and next block’s task within a block. Task-label permutation tests reassigned the six task labels of every session among themselves, independently for each session, so that all blocks of a task kept one shared label (across-session analyses, Figures 4 and 5 and Supplementary Figure 4). For the correlations with reward amount, saccade reaction time and hit rate (Figure 6), the same permutation was applied to all sessions. Area-label and monkey-label permutation tests permuted the area or monkey labels of sessions (Figure 5A and B, Supplementary Figure 5, and FEF versus dlPFC with 100 neurons per session). Sign-flip tests randomly flipped the signs of a per-session paired difference between two measures (Supplementary Figure 2A and C). Time-shuffle tests permuted the time positions of the blocks within a session (Supplementary Figure 2B). All tests were two-sided, except the one-sided test of the gain from adding a component (Figure 3A and C). When the observed value was more extreme than every permutation, p took its minimum value, 1/(number of permutations + 1), and was reported as p < 0.001 for 1,000 permutations and p < 0.0001 for 10,000 permutations. Unless stated otherwise, values are given as mean ± SD across sessions, to show the variability across sessions, and statistical significance was assessed with the permutation tests described above.

Unless stated otherwise, group-level tests over sessions used the mean of the per-session statistic. Its null distribution was built by drawing, for each session, one value at random from that session’s own permutation distribution and averaging across sessions. This was repeated 10,000 times, and p-value was the fraction of null means at least as far from the mean of the null distribution as the observed mean, in either direction (observed value included). In the across-session analyses, each session contributes to many comparisons, so the task labels of every session were permuted independently. For the similarity of session pairs (Supplementary Figure 4), the mean similarity was recomputed for 10,000 such permutations. For the cross-session decoding (Figures 4B, 5A and 5B), the whole analysis was repeated for 1,000 such permutations.

Same-area versus other-area training (Figure 5A) and same-monkey versus other-monkey training (Figure 5B) were compared by permuting the area or monkey labels of the sessions. The number of sessions per group was kept, and the size-matched training sets were rebuilt from the permuted labels. The mean difference in accuracy over the test sessions was then compared with its permutation distribution. For Figure 5A, area labels were permuted within each monkey, using all C(8, 3) = 56 relabelings in Monkey F and 1,000 random relabelings in Monkey H. For Figure 5B, monkey labels were permuted across all 32 sessions of both monkeys, using 1,000 random relabelings.

## Author contributions

JP designed the experiment, analyzed the data, and wrote the manuscript. HP and CH conducted the experiment and collected the data. LS helped design the experiment and write the manuscript.

## Acknowledgments

We thank H. Song, X. Wei, and T. Elston for helpful discussion and feedback regarding this work and J. Tucker for helping with technical issues and collecting data.

## Supplementary Information

**Supplementary Figure 1.**
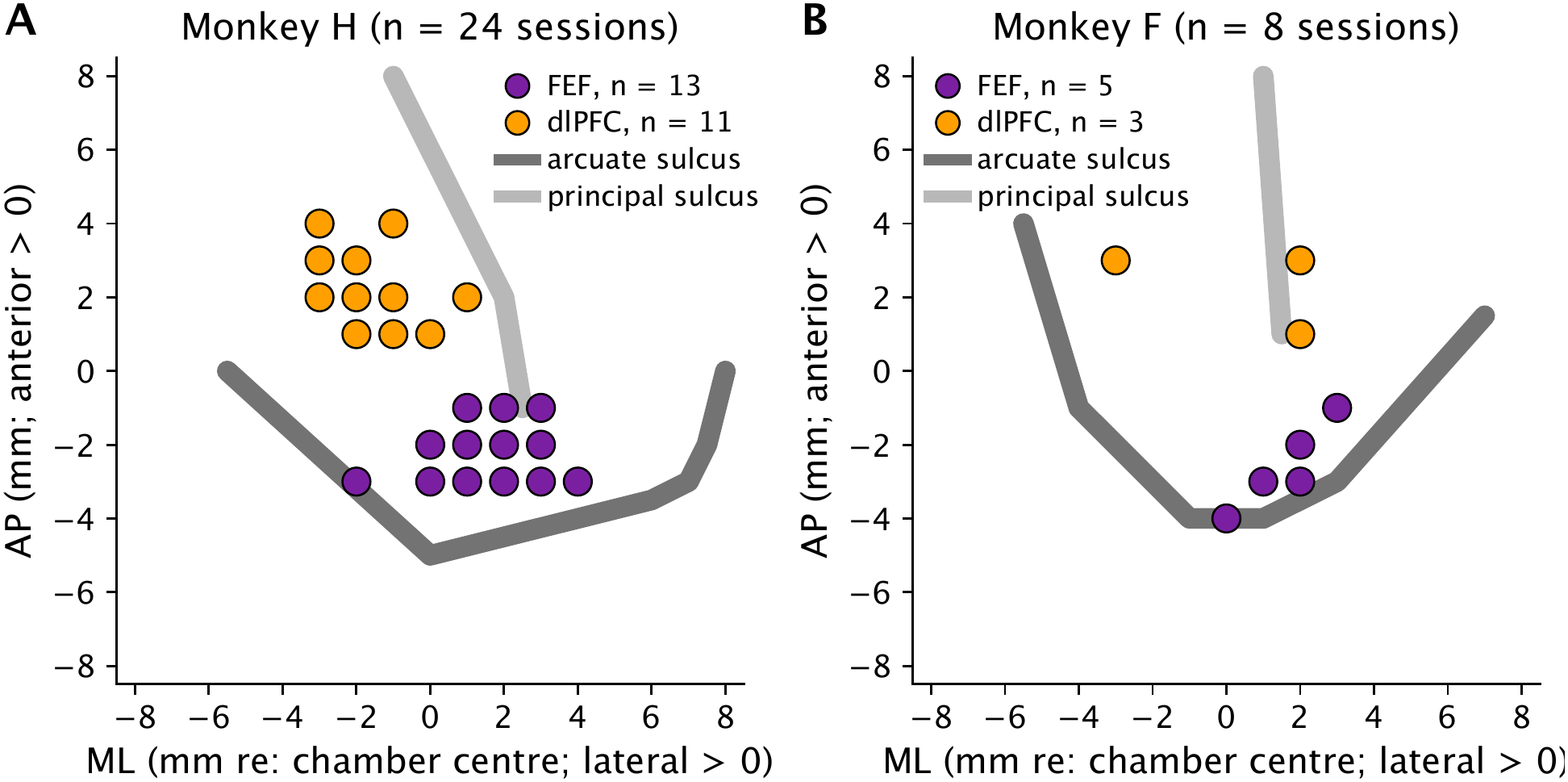
Recording sites (A–B) Recording sites of all sessions of Monkey H (A, 13 FEF and 11 dlPFC sessions) and Monkey F (B, 5 FEF and 3 dlPFC sessions) in chamber coordinates (1-mm grid, positive values lateral and anterior). Every session was recorded from different recording sites. Gray lines, sulcus outlines from the chamber maps (dark for the arcuate sulcus and light for the principal sulcus).

**Supplementary Figure 2.**
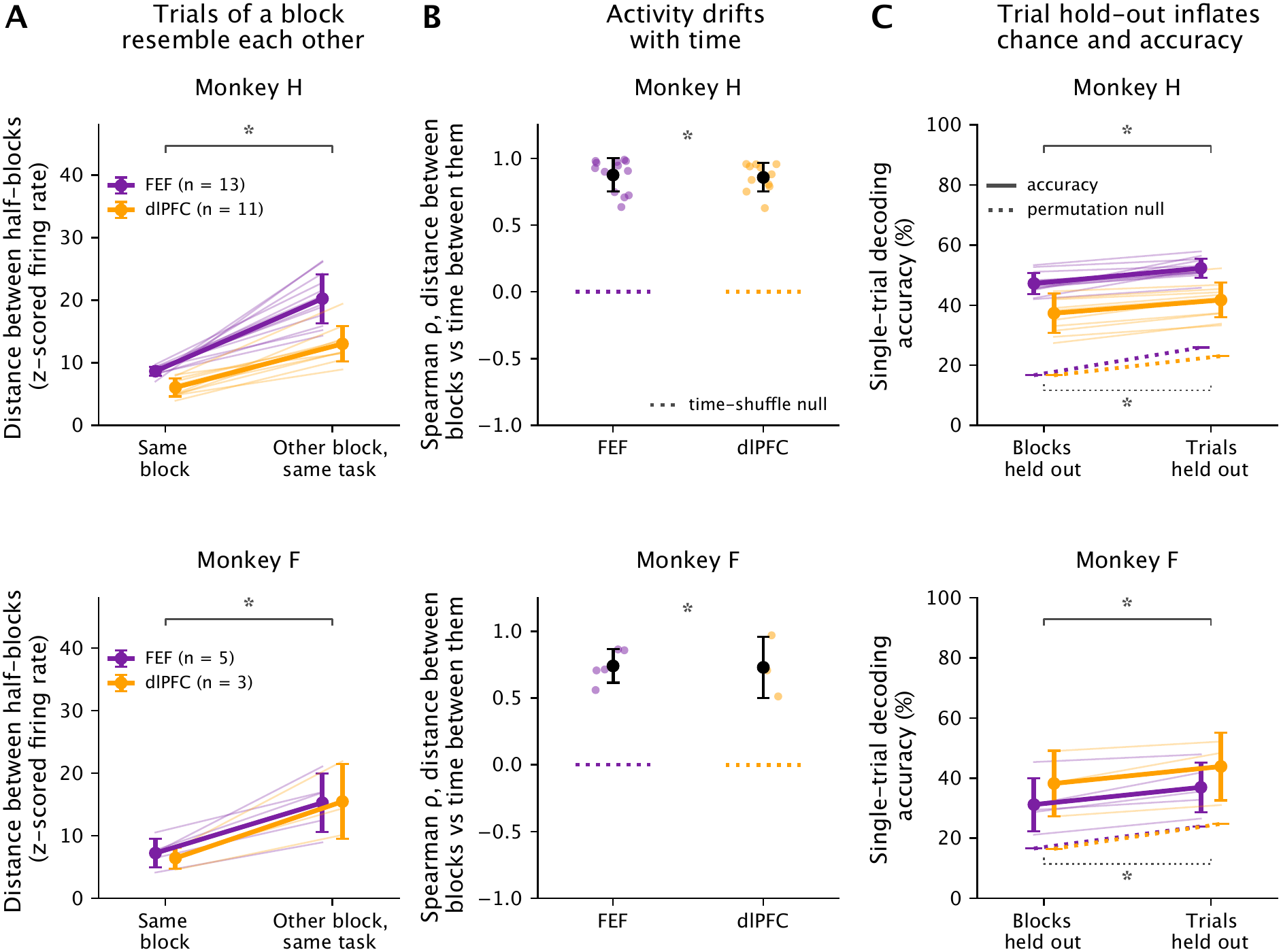
Slow drift makes trials within a block alike. **(A)** Euclidean distance across all neurons between two halves of the successful trials of a block (firing rates z-scored per neuron across trials, then averaged within each half): the two halves of the same block (left) or a half of one block and a half of another block of the same task (right). Thick lines and error bars: mean ± SD across the sessions of each area (purple for FEF and orange for dlPFC). Thin lines: sessions. The two halves of a block were closer to each other than to halves of other blocks of the same task in every session (7.5 ± 1.7 versus 16.9 ± 5.0 in Monkey H and 6.9 ± 2.0 versus 15.3 ± 4.8 in Monkey F, mean ± SD across sessions). Bracket: sign-flip test of the paired differences across the sessions of each monkey, FEF and dlPFC pooled (p < 0.0001 in Monkey H and p = 0.008 in Monkey F). Asterisk, p < 0.05. **(B)** Spearman correlation between the distance of two blocks of the same task (as in A, from block means) and the time between them (in each session). Dots: sessions. Black: mean ± SD across the sessions of each area. Dotted lines: mean of the time-shuffle null, in which the time positions of the blocks were shuffled within each session (1,000 shuffles). Blocks farther apart in time differed more in every session (ρ = 0.87 ± 0.11 in Monkey H and 0.74 ± 0.16 in Monkey F, versus 0.00 ± 0.03 under the null, mean ± SD across sessions). Asterisk: mean ρ of the sessions of each monkey, FEF and dlPFC pooled, against the time-shuffle null (p < 0.0001 in both monkeys). **(C)** Single-trial decoding accuracy (first four components) with whole blocks or single trials held out. Solid lines: accuracy. Thick lines and error bars: mean ± SD across the sessions of each area. Thin lines: sessions. Dotted lines: mean of the block-label permutation null (50 permutations per session). Holding out single trials raised the null from chance (16.7%) to 24.6 ± 3.4% in Monkey H and 24.2 ± 3.5% in Monkey F (mean ± SD across sessions), because the other trials of a held-out trial’s block stayed in the training set with the same permuted label. Accuracy also rose, by 4.8 ± 2.5 percentage points in Monkey H and 5.7 ± 3.1 in Monkey F (mean ± SD), in every session. Brackets: sign-flip tests of the paired differences for accuracy (top) and null (bottom) across the sessions of each monkey, FEF and dlPFC pooled, p < 0.0001 in Monkey H and p = 0.008 in Monkey F. Asterisk, p < 0.05.

**Supplementary Figure 3.**
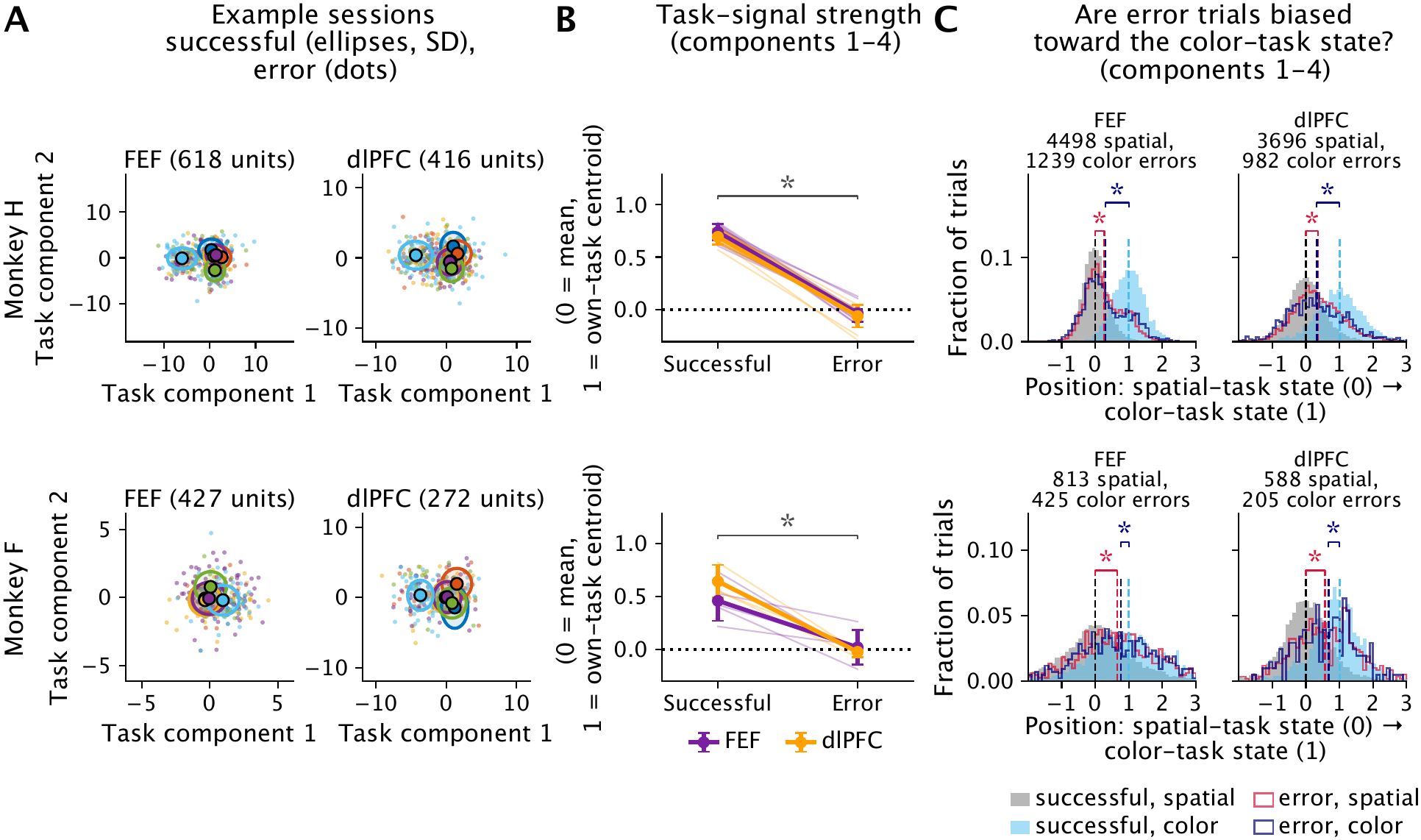
Error trials lack the task-specific population state. **(A)** Held-out successful and error trials of the example sessions of Figure 2B on the first 2 task components. Ellipses: mean ± SD of the successful trials of each task. Small dots: error trials. **(B)** Task-signal strength of held-out trials: the mean position of the successful and of the error trials of each session on the axis from the mean of all 6 tasks (0) to the centroid of the trial’s own task (1) in the task subspace defined by the first 4 components. Thin lines: sessions. Thick lines and error bars: mean ± SD across the sessions of each area (purple for FEF and orange for dlPFC). Bracket: successful versus error trials, trial-label permutation test of success and error within each session, sessions combined. Asterisk, p < 0.05. **(C)** Position of held-out trials on the axis from the mean of the successful spatial-task trials (0) to the centroid of the successful color-task trials (1) in the task subspace defined by the first 4 components. Pooled over the sessions of each area. Histograms show the fraction of trials of each group (gray: successful spatial-task trials, light blue: successful color-task trials, red outline: error trials of the spatial tasks, navy outline: error trials of the color task). Dashed lines: group means. Brackets: trial-label permutation tests of success and error within each session for the shift of spatial-task error trials toward the color-task state (red) and of color-task error trials toward the spatial-task state (navy). Asterisk: p < 0.05.

**Supplementary Figure 4.**
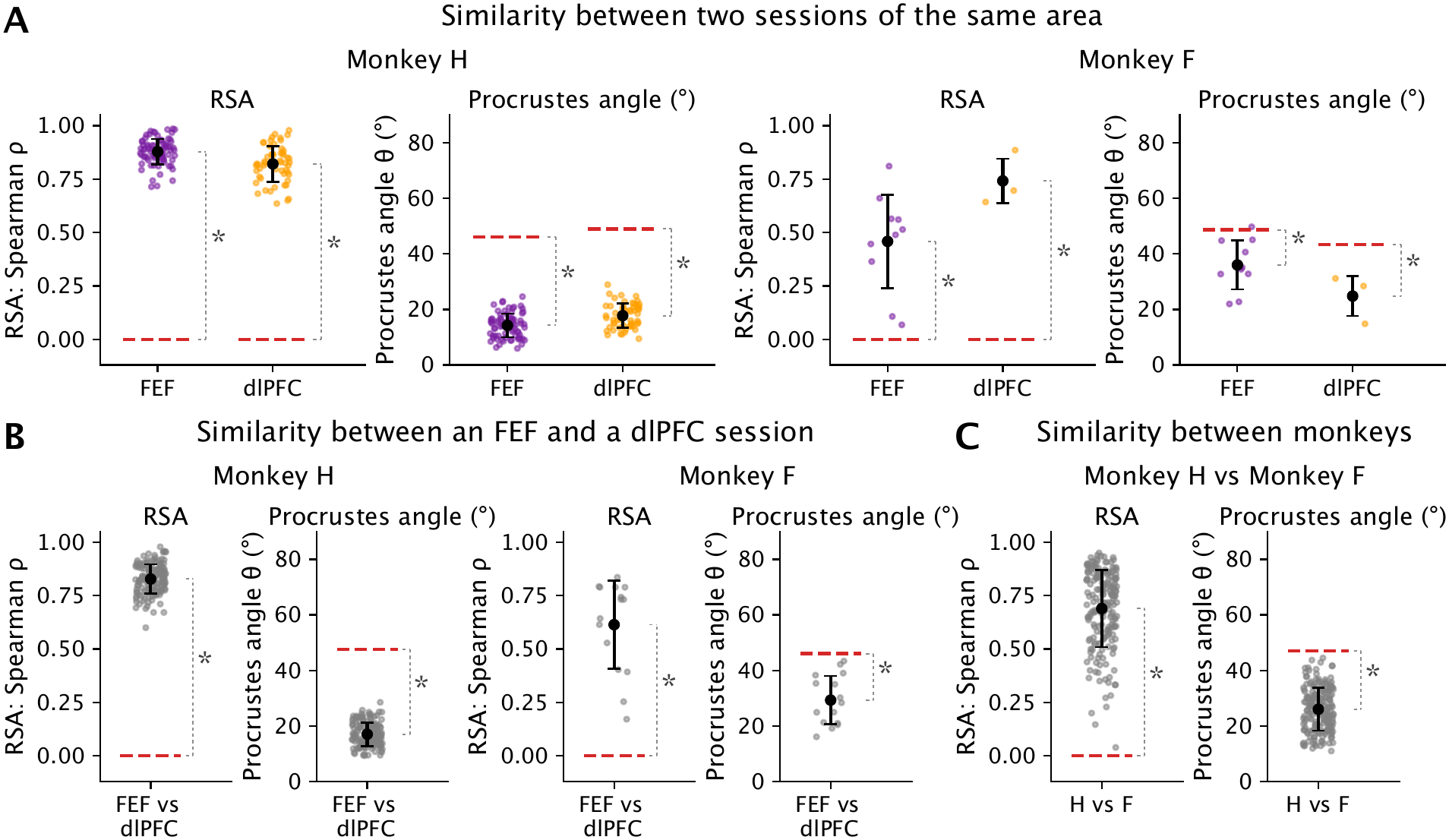
Similarity of the task geometry between sessions, areas and monkeys. **(A)** Pairwise similarity of the task geometries of 2 sessions of the same area, first 4 components: RSA (Spearman correlation between the 15 inter-centroid distances, left, where perfect alignment approaches 1) and Procrustes angle after optimal alignment (right: perfect alignment approaches 0). Small colored dots: session pairs. Black dot and error bar: mean ± SD. Red dashed lines: null (mean similarity of a pair over all 720 relabelings of the tasks of one session, averaged over pairs). Asterisks: task-label permutation test of the 6 tasks within every session (10,000 permutations, p < 0.05). **(B)** Similarity of the task geometries of an FEF session and a dlPFC session, for every within-animal FEF–dlPFC session pair (143 pairs in Monkey H and 15 in Monkey F), first 4 components: RSA (Spearman correlation between the 15 inter-centroid distances, left) and Procrustes angle after optimal alignment (right). Small dots: session pairs. Black dot and error bar: mean ± SD. Red dashed lines: null similarity. Asterisks: p < 0.05 in the task-label permutation test of the 6 tasks within every session (10,000 permutations). **(C)** Similarity of the task geometries of a Monkey H session and a Monkey F session, for every Monkey H–Monkey F session pair (192 pairs), first 4 components: RSA (left) and Procrustes angle (right). Small dots: session pairs. Black dot and error bar: mean ± SD. Red dashed lines and asterisks are as in B.

**Supplementary Figure 5.**
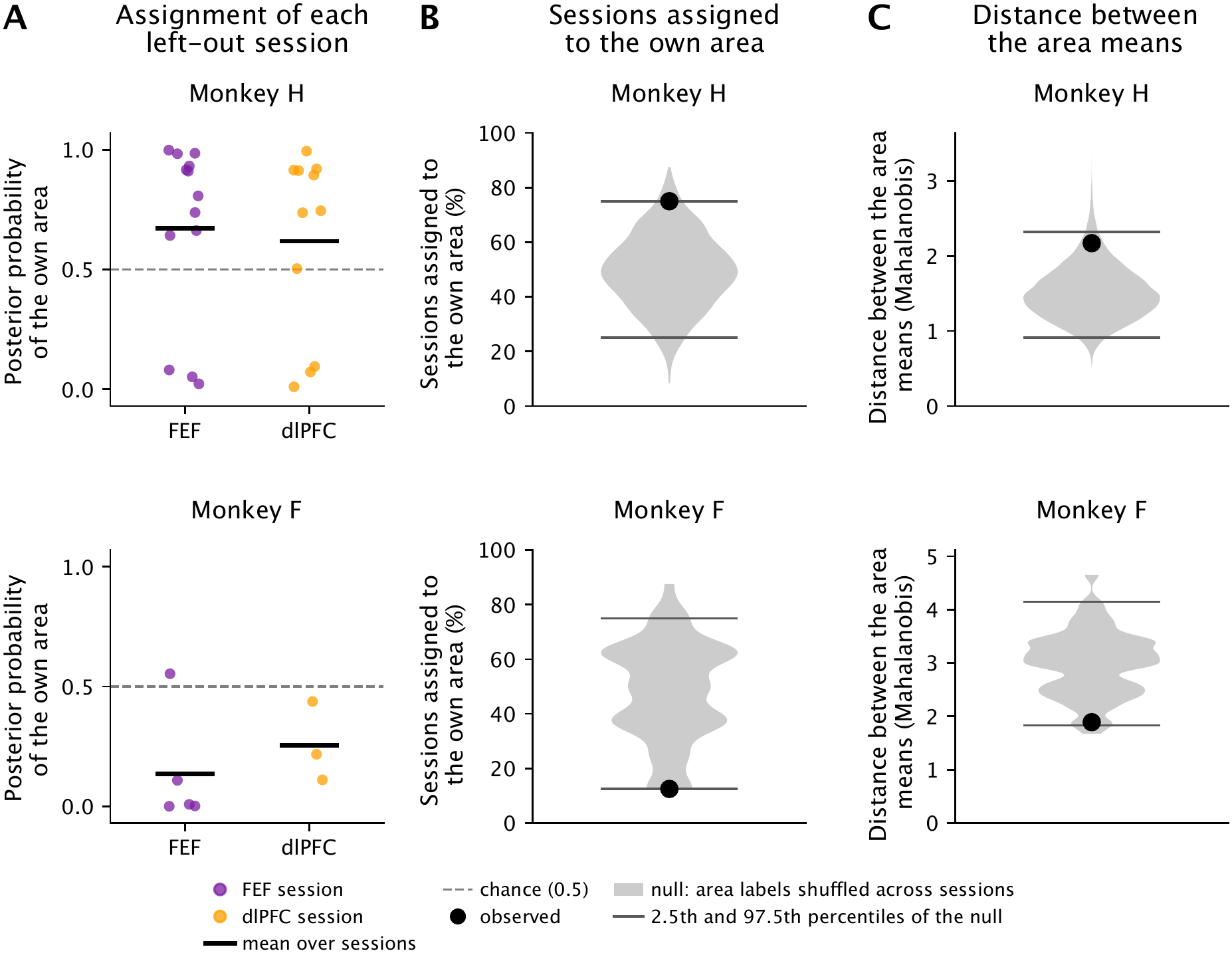
Area identity of the task geometry. **(A)** Assignment of each left-out FEF or dlPFC session to an area, first 4 components, in Monkey H (top, 13 FEF and 11 dlPFC sessions) and Monkey F (bottom, 5 and 3 sessions). The geometry of a session was its 15 cross-validated squared distances between task centroids, divided by their mean. Each session was assigned to the area whose mean geometry, computed from the other sessions, gave the higher likelihood. Dots: posterior probability of the session’s own area. Black lines: mean over sessions. Dashed line: chance. **(B)** Percentage of sessions assigned to their own area. Black dot: observed. Gray violin: null distribution when the area labels were shuffled across sessions and the whole procedure was repeated (1,000 shuffles). Dark ticks: the 2.5th and 97.5th percentiles of the null. **(C)** Mahalanobis distance between the mean geometries of FEF and dlPFC (black dot) and its null distribution (gray violin, 10,000 shuffles). Dark ticks as in B.

---

**Supplementary Table 1.** Similarity of task geometry between sessions of the same area.

|  | **Monkey H<br>FEF** | **Monkey H<br>dlPFC** | **Monkey F<br>FEF** | **Monkey F<br>dlPFC** |
|---|---|---|---|---|
| RSA ρ | 0.88 | 0.82 | 0.46 | 0.74 |
| RSA, permutation p | < 0.0001 | < 0.0001 | 0.001 | 0.0006 |
| RSA, session pairs above null | 94% | 84% | 10% | 67% |
| Procrustes angle | 14° | 18° | 36° | 25° |
| Procrustes angle, task-permuted | 46° | 49° | 49° | 43° |
| Cosine similarity, permutation p | < 0.0001 | < 0.0001 | 0.0005 | 0.0003 |
| Cosine similarity, session pairs above null | 100% | 98% | 20% | 67% |

RSA ρ: Spearman correlation between the pairwise centroid distances of two sessions of the same area. Procrustes angle: angle between the centroid configurations of two sessions after centering, normalizing to unit size and optimal alignment by rotation and reflection. Task-permuted: the same angle for geometries with the 6 task labels permuted within every session. Permutation p: group-level task-label permutation test of the 6 tasks within every session (Supplementary Figure 4A). Session pairs above null: percentage of individual session pairs whose similarity exceeded their own task-label permutation null (p < 0.05).

**Supplementary Table 2.** Cross-session decoding of task blocks (first 4 components)

|  | **Monkey H<br>FEF** | **Monkey H<br>dlPFC** | **Monkey F<br>FEF** | **Monkey F<br>dlPFC** |
|---|---|---|---|---|
| Leave-one-session-out, components 1–4 (Figure 4B) | 66 ± 16% | 62 ± 16% | 24 ± 14% | 42 ± 5% |
| Components 1–4, p | < 0.001 | < 0.001 | 0.055 | 0.003 |
| Components 1–4, sessions above null | 12 of 13 | 10 of 11 | 1 of 5 | 3 of 3 |
| Component 1 | 46 ± 9% | 39 ± 5% | 28 ± 7% | 35 ± 4% |
| Gain from 1 to 4 components, p | < 0.001 | < 0.001 | 0.29 | 0.30 |
| Change from 4 to 5 components (percentage points) | −1.1 ± 3.2 | −6.9 ± 9.1 | 1.0 ± 4.8 | −7.5 ± 9.2 |
| Change from 4 to 5 components, p | 0.68 | 0.011 | 0.68 | 0.058 |
| Component 5 alone | 18 ± 3% | 16 ± 4% | 17 ± 3% | 17 ± 4% |
| Component 5 alone, p | 0.36 | 0.80 | 0.97 | 0.91 |
| Same-area training (Figure 5A) | 67 ± 15% | 62 ± 16% | 21 ± 6% | 42 ± 5% |
| Same-area training, p | < 0.001 | < 0.001 | 0.054 | 0.003 |
| Other-area training | 62 ± 11% | 59 ± 15% | 34 ± 16% | 37 ± 5% |
| Other-area training, p | < 0.001 | < 0.001 | < 0.001 | < 0.001 |
| Same versus other area, p | 0.28 | 0.19 | 0.018 | 0.61 |
| Same-monkey training (Figure 5B) | 64 ± 12% | 59 ± 15% | 29 ± 14% | 45 ± 12% |
| Same-monkey training, p | < 0.001 | < 0.001 | 0.002 | < 0.001 |
| Other-monkey training | 62 ± 16% | 47 ± 13% | 29 ± 13% | 46 ± 4% |
| Other-monkey training, p | < 0.001 | < 0.001 | < 0.001 | < 0.001 |
| Same versus other monkey, p | 0.77 | 0.027 | 0.86 | 0.73 |

Accuracy: mean ± SD across test sessions, chance 16.7%. p: task-label permutation test of the 6 tasks within every session, with the whole analysis repeated (1,000 permutations), except for same versus other area or monkey (area-label or monkey-label permutation test of the sessions). Leave-one-session-out: all other sessions of the test area, aligned in components 1 to k for each number of components k (Figure 4B). Same-area and other-area training: 10 training sessions in Monkey H and 2 in Monkey F (Figure 5A). Same-monkey and other-monkey training: 7 training sessions (Figure 5B).

**Supplementary Table 3.** Similarity of task geometry between areas and between monkeys.

|  | **Monkey H<br>FEF–dlPFC** | **Monkey F<br>FEF–dlPFC** | **Monkey H–<br>Monkey F** |
|---|---|---|---|
| Session pairs | 143 | 15 | 192 |
| RSA ρ | 0.83 | 0.61 | 0.69 |
| RSA, permutation p | < 0.0001 | < 0.0001 | < 0.0001 |
| RSA, session pairs above null | 88% | 47% | 54% |
| Procrustes angle | 17° | 29° | 26° |
| Procrustes angle, task-permuted | 48° | 46° | 47° |
| Cosine similarity, permutation p | < 0.0001 | < 0.0001 | < 0.0001 |
| Cosine similarity, session pairs above null | 99% | 40% | 53% |

As in Supplementary Table 1, for the pairs of an FEF and a dlPFC session of the same monkey (Supplementary Figure 4B) and for the pairs of a Monkey H and a Monkey F session (Supplementary Figure 4C).

